# A single intracellular protein governs the critical transition from an individual to a coordinated population response during quorum sensing: Origins of primordial language

**DOI:** 10.1101/074369

**Authors:** Celina Vila-Sanjurjo, Christoph Engwer, Xiaofei Qin, Lea Hembach, Tania Verdía-Cotelo, Carmen Remuñán-López, Antón Vila-Sanjurjo, Francisco M. Goycoolea

## Abstract

Quorum sensing (QS) explains a type of bacterial cell-cell communication mediated by exocellular compounds that act as autoinducers (AIs). As such, QS can be considered the most primordial form of language. QS has profound implications for the control of many important traits (*e.g.* biofilm formation, secretion of virulence factors, etc.). Conceptually, the QS response can be split into its “listening” and “speaking” components, *i.e.* the power to sense AI levels *vs.* the ability to synthesize and release these molecules. By explaining the cell-density dependence of QS behavior as the consequence of the system’s arrival to a threshold AI concentration, models of QS have traditionally assumed a salient role for the “QS speaking” module during bacterial cell-to-cell communication. In this paper, we have provided evidence that challenges this AI-centered view of QS and establishes LuxR-like activators at the center of QS. Our observation that highly coordinated, cell-density dependent responses can occur in the absence of AI production, implies that the ability to launch such responses is engrained within the “QS listening” module. Our data indicates that once a critical threshold of intracellular activator monomers in complex with AI is reached, a highly orchestrated QS response ensues. While displaying a clear cell-density dependence, such response does not strictly require the sensing of population levels by individual cells. We additionally show, both *in vivo* and *in silico*, that despite their synchronous nature, QS responses do not require that all the cells in the population participate in the response. Central to our analysis was the discovery that percolation theory (PT) can be used to mathematically describe QS responses. While groundbreaking, our results are in agreement with and integrate the latest conclusions reached in the field. We explain for the first time, the cell-density-dependent synchronicity of QS responses as the function of a single protein, the LuxR-like activator, capable of coordinating the temporal response of a population of cells in the absence of cell-to-cell communication. Being QS the most primordial form of speech, our results have important implications for the evolution of language in its ancient chemical form.

**Abbreviations:**

3D: three dimensional
*a*_*c*_: wthreshold intracellular concentration of activator molecules
AHL: acyl-homoserine lactone
AHL _*fisch*_: N-(3-oxohexanoyl)-L-homoserine lactone
AHL _*viol*_: N-hexanoyl-DL-homoserine-lactone
AI: autoinducer
a.u: arbitrary units
BMB: bromophenol blue
CA: trans-cinnamaldehyde
Fl: fluorescence intensity
FI/OD600: density-normalized fluorescence intensity
GFP: green fluorescent protein
M_*w*_: molecular weight
PT: percolation theory
QS: quorum sensing
t_*c*_: percolation critical time
wt: wild type

---

Quorum sensing (QS) explains a type of bacterial cell-to-cell communication mediated by exocellular chemical compounds that act as autoinducers (AIs). Since QS can be considered the most primordial form of language [1–6], understanding its chemical basis is required in order to gain a proper perspective on the evolution of speech. Conceptually, the QS response can be split into its “listening” and “speaking” components, *i.e.* the power to sense AI levels *vs.* the ability to synthesize and release these molecules. Traditionally, an AI centered model has been used to explain QS’s dependence on cell density. In such a model the accumulation of AIs up to a threshold level during cell growth is what determines the timing of the response [7].

Given that cell-to-cell communication by QS has profound implications in control of many important bacterial traits (*e.g.* biofilm formation, secretion of virulence factors, etc.), QS has been a target for multidisciplinary research in the past few years. Indeed, not only has it attracted the attention of the Microbiology community but also of researchers in the fields of Systems and Synthetic Biology [8, 9]. As a result, efforts towards the construction of mathematical models that can accurately describe QS responses have gained considerable traction. A recurrent feature revealed by many of these mathematical models is the importance of gene expression noise in shaping the QS response [8, 10–19]. Despite all these advances, we still do not understand how individual cellular responses translate into the cell-density-dependent synchronicity observed at the population level. Understanding this aspect of QS is key for the development of an accurate model of bacterial cell-to-cell communication.

Since the cell-density-dependent behavior of the whole system arises from the complex interplay of the microscopic, biochemical parameters (*e.g.*, species concentrations, reaction rates, binding constants, gene expression rates, and so on), attempting to simulate the population behavior based on these parameters appears as an insurmountable task. Not surprisingly, approaches towards the simplification of mechanistic models of QS have been proposed [16]. Percolation theory (PT) is a general mathematical formalism that describes connectivity and transport in geometrically complex systems. Specifically, PT describes the behavior of a system in the vicinity of a critical threshold above which, an infinite cluster of interconnected objects spans a given network system. Notably, the mathematical simplicity of PT makes it a highly convenient tool to explain a diversity of complex physical and biological processes that occur as a result of critical phenomena [20–23]. Here, we show, for the first time, that PT can be used to conveniently describe experimental QS responses and that this feature can be used to validate the results of computer simulations connecting single-cell microscopic events to the phenotypic behavior of the population.

By combining *in vivo* measurements with *in silico* simulations, we have elucidated how gene expression noise in individual cells can be integrated into highly orchestrated responses at the population level. First, we use a cellular model of “QS listening” to show that the generation of a density-dependent, highly synchronized response can occur in the absence of cell-to-cell communication. Second, we show that PT can be conveniently used to mathematically approximate QS responses. Third, we create a simple algorithm capable of recreating most of the features displayed by our cellular models of “QS listening”. Finally, by combining our *in vivo* and *in silico* data, we arrive at a model of QS in which cell-density-dependent synchronicity is solely driven by the stochasticity associated with the expression of the QS activator, acting within the constraints imposed by growth dynamics.

## Results

### Cell-density-dependent synchronicity occurs in the absence of AHL production

It has been amply demonstrated that the production of the pigment violacein by *Chromobacterium violaceum* is regulated by QS [24]. The *C. violaceum* QS apparatus consists of the LuxR/I-like proteins CviR and CviI [25]. The *C. violaceum* strain CV026 is a well-known QS biosensor that has lost the ability to produce the AI N-hexanoyl-L homoserine lactone (C6HSL, AHL _*viol*_) due to a double Tn5 transposon insertion mapped to the *cviI* gene, which codes for the AHL _*viol*_ synthase (CviI), and to a putative repressor locus [24]. Despite these insertions, *C. violaceum* CV026 can still respond to the presence of AHL _*viol*_ and release violacein [26]. Since this biosensor cannot produce AHLs by itself, individual cells cannot receive information about any AHL-induced responses occurring in their neighborhood, except for their own. A direct consequence of this is that the cells should not be able to sense population levels via the CviR/CviI system. Hence, this “mute” *C. violaceum* strain constitutes an exquisite model to study “QS listening”. When *C. violaceum* CV026 cells are plated in the presence of sufficiently high concentrations of AHL _*viol*_, a concentric ring of violacein is produced around the AHL _*viol*_ source after prolonged incubation (Figure 1A). The fact that no violacein is produced outside the ring attests to the well-known inability of *C. violaceum* CV026 to synthesize AHL (Figure 1A) [24, 27, 28]. In Figure 1B we plotted violacein intensity across the ring against distance from the AHL _*viol*_ source, at three of the AHL _*viol*_ concentrations shown in Figure 1A. A saturation plateau, spanning almost the whole width of the ring, is followed by a region where violacein intensity decays in a sigmoidal fashion. To better understand the shape of the underlying gradient of AHL _*viol*_ we modeled its diffusion in agar by using bromophenol blue (BMB) as a surrogate, diffusible probe with low *Mw* (see Materials and Methods). Figure 1C shows that the time-dependent diffusion of BMB into agar follows a monotonic exponential decay function (see also Figure S1). This result is in agreement with the results of Anbazhagan *et al.* [29], who found a similar relationship by measuring the dependence of AHL _*viol*_-induced rings on the concentration of the inducer. Clearly, the sigmoidal character of violacein intensity across the ring (*c.f.* Figures 1B and C), indicates that the kinetics of the QS response launched by *C. violaceum* CV026 cells does not mirror the underlying AHL _*viol*_ diffusion gradient.

**Figure 1.**
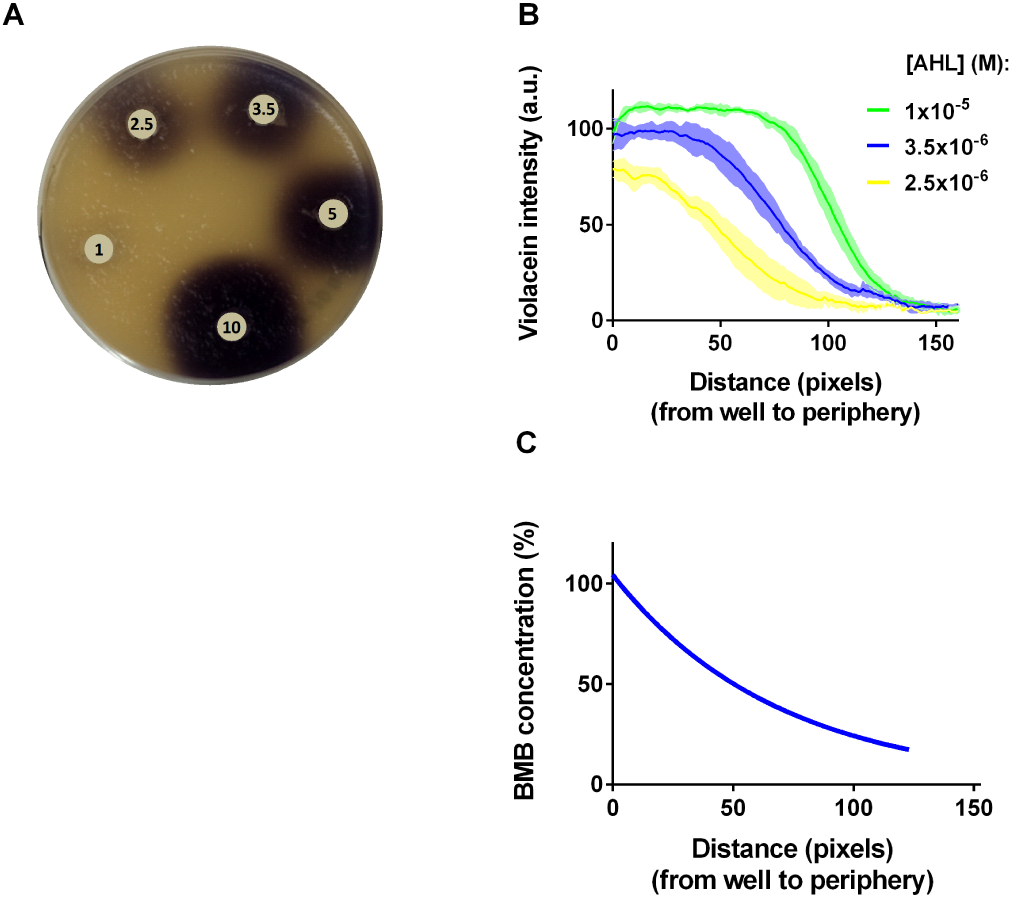
End-point QS response of the C. violaceum biosensor. A. Plate assay showing the formation of violacein rings by *C. violaceum* CV026 induced with varying AHL _*viol*_ concentrations (numbers shown in wells represent AHL _*viol*_ concentrations in the µM range). B. Variation in violacein intensity within the rings represented in Figure 1A as a function of the distance from the edge of the well. X-Axis: distance from the edge of the well measured in pixels. Y-axis: Violacein intensity measured in arbitrary units. Green, 10µM AHL _*viol*_; blue 3.5 µM AHL _*viol*_; and yellow 2.5 µM AHL _*viol*_. Solid lines represent the mean. Light-colored, shaded areas represent the standard deviation from three biological replicates. C. Modeled concentration of BMB as a function of distance after diffusion in agar for 48 h. X-Axis: distance from the edge of the well measured in pixels. Y-axis: relative concentration of BMB in %.

To better understand the temporal kinetics of violacein accumulation, we performed time-lapse analysis of *C. violaceum* CV026 cells in the presence of external AHL _*viol*_. The results are shown in Figure 2A and Movie S1 (note that the experiment has been performed at room temperature to achieve increased time resolution, see Materials and Methods). Under these conditions, a transition from clear to opaque agar at ∼32 h, caused by cell growth, is followed by the sudden appearance of a violet hue at ∼48 h. Violacein keeps accumulating until the end of the experiment (80 h) (Figure 2A). At no point in time, violacein accumulation appears to be proportional to the underlying gradient of AHL _*viol*_ concentration (modeled in Figure 1C), but rather resembles an all-or-none response (Figure 2A and Movie S1).

**Figure 2.**
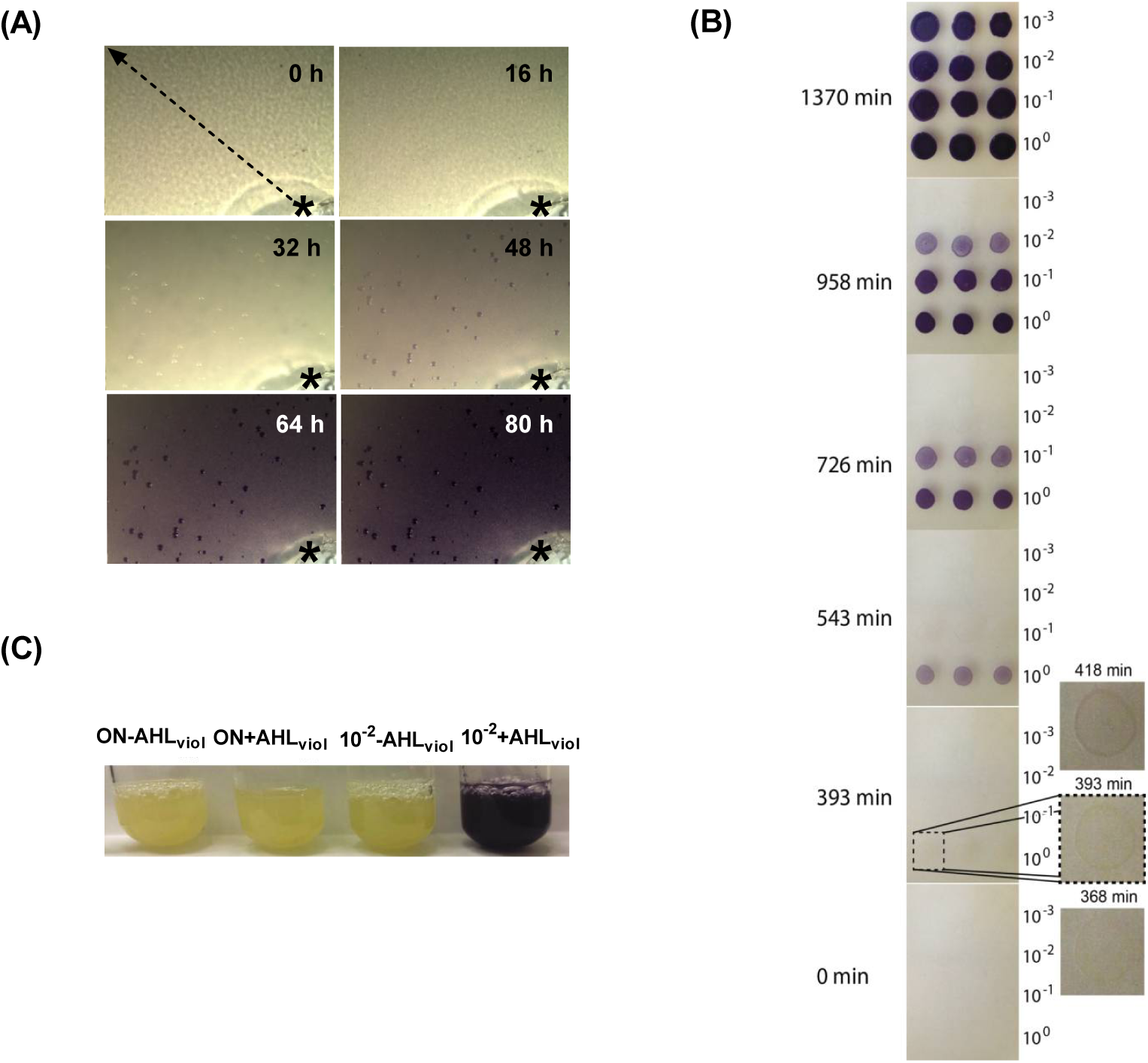
Time-lapse QS response of the C. violaceum biosensor. A. Violacein production at 0, 16, 32, 48, 64, and 80 h of incubation of *C. violaceum* CV026 cells in the presence of AHL _*viol*_. The position of the AHL well, containing 50 µL of an aqueous 10-µM solution of AHL _*viol*_ at the beginning of the assay, is indicated by an asterisk. Violacein intensity was measured along the direction marked with the arrowed line shown at time 0. B. Violacein production *vs.* cell density. The panels show the accumulation of violacein relative to time after plating 2-µL drops of 100, 10^−1^, 10^−2^, and 10^−3^ serial dilutions of a stationary-phase culture of *C. violaceum* CV026 onto AHL _*viol*_-containing LB-agar. The time after plating and the dilution are indicated. Note that each frame beyond 0 min, except for the last one at 1370 min, corresponds to a turning point at which violacein becomes visible for one of the serial dilutions. The inset on the right shows an enlarged view of one of the drops at the 393-min frame and a similar view of the same area in the previous and subsequent frames. C. Lack of pigment production by stationary-phase cultures of *C. violaceum* CV026 cells after addition of AHL _*viol*_. A 10^−2^ dilution of the same culture was used as a positive control.

It has been reported that violacein accumulates in *C. violaceum* upon entry in stationary phase [30, 31]. High-resolution spectroscopy measurements of violacein production by wild-type (wt) cells revealed the detection of the pigment ∼6 h after plating stationary-phase cells on fresh agar [28]. When we plate the *C. violaceum* CV026 strain under the similar conditions and in the presence of external AHL _*viol*_, comparable delays of ∼6 h are observed (Figure 2B, 393-min time point). This indicates that despite the lack of the “QS speaking” module in this strain, the QS response follows comparable kinetics in both strains. Indeed, the experiment of Figure 2B shows that cell-density-dependence, the hallmark of QS, is maintained in the “mute” strain. By plating serial dilutions of *C. violaceum* CV026 stationary-phase cells in the presence of AHL _*viol*_, we observed that visible levels of violacein appeared at drastically different times (Figure 2B, Movie S2). The frames shown in the figure are the first time points at which violacein accumulation became visible as a faint circle around the plated cells (see inset in Figure 2B comparing the cells at 368 min, the frame before the appearance of visible violacein; 393 min, the first frame in which violacein becomes apparent; and 418min, the frame after the appearance of visible violacein). The closer the cell density was to its original value in the stationary-phase culture, the earlier the appearance of visible violacein (Figure 2B and Movie S2). At the same time, these results clearly rule out the alternative explanation, namely that violacein biosynthesis could be the rate-limiting step of the observed response. Strikingly, the addition of AHL _*viol*_ to stationary-phase cultures resulted in no violacein production (Figure 2C). These results provide diagnostic evidence that even in the absence of cell-to-cell communication, due to the lack of the QS “speaking” module, growing *C. violaceum* CV026 are capable of detecting cell density. The cells do not merely respond to the presence of external AI, but wait until they reach a certain density and only then, they respond. Since plenty of AI is present from time zero in the system, this observation is in stark contrast to the tenets of the standard QS model, namely the requirement of a steady buildup of AI up to a threshold level. Notably, this ability to respond to cell-density is lost in non-growing *C. violaceum* CV026 cells.

Taken together, these results demonstrate that even in the absence of cell-to-cell communication via the CviR/CviI system, *C. violaceum* CV026 cells can elicit a highly concerted, delayed response in the presence of externally added AHL _*viol*_. Strikingly, its dependence on high cell density levels makes this response indistinguishable from that of a standard QS event in which both the “listening” and “speaking” modules are present.

### The macroscopically observed QS response is elicited by a minority of the cells

Figure 3A shows a view of a violacein ring at higher magnification. A wide area of active violacein deposition, represented by frames 1–7, is followed by a region of rapid decay, frames 8–10. While most cells form a lawn within the thin layer of soft agar on top of the plate, interspersed large colonies are also visible on the surface of the agar (Figure 3A). Figures 3A and B also show the existence of discrete patches of violacein deposition within the ring, suggesting that not all the cells within the cell lawn were contributing to pigment deposition. To dwell further into this issue, we subjected violacein-producing, stationary-phase cultures of *C. violaceum* CV026 to brightfield microscopy under high magnification. Strikingly, we observed that at the microscopic level, violacein production was restricted to scattered patches of cells, each one containing a small number violacein-producing units (Figure 3C, Movie S3). Therefore, exposure of *C. violaceum* CV026 to AHL _*viol*_ results in an apparently stochastic system of violacein-producing cells.

**Figure 3.**
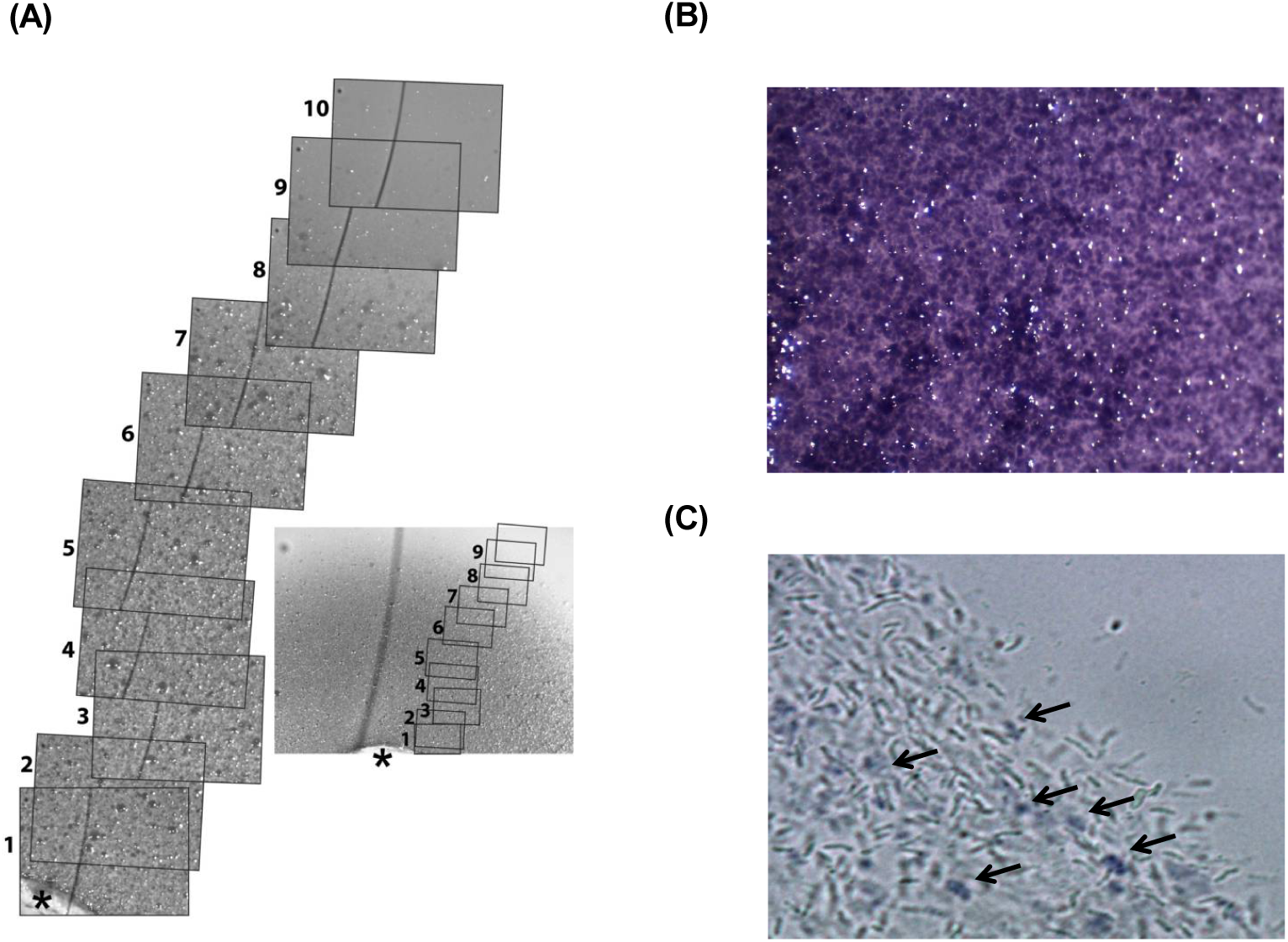
Time-lapse analysis of violacein production relative to cell density. A. View of the violacein ring at low magnification after 2 days of growth at 30*◦*C. The position of the AHL well, containing 50 µL of an aqueous 10-µM solution of AHL _*viol*_ at the beginning of the assay, is indicated by an asterisk. The relative position of the individual frames is shown on the lower magnification view. The smooth black line, present in all frames, is due to a defect on the camera’s microsensor. B. Low-magnification image of the violacein ring of a bioassay plate, showing purple and clear patches. C. Bright-field image of a mucoid material from a stationary-phase, liquid culture of *C. violaceum* CV026 (see Materials and Methods). The position of purple cells is indicated with arrows.

### The “QS listening” ability of *Chromobacterium violaceum* displays “percolation kinetics”

The abrupt rise in violacein intensity within a narrow concentration range observed at the periphery of the ring, together with the apparent synchronicity of this process across the ring, strongly suggested that the underlying kinetics could be those of a critical phenomenon and, as such, could be approximated by percolation theory (PT) [21]. To test this hypothesis, we fitted the evolution of violacein deposition by *C. violaceum* CV026 (data from Figure 2A) over time to Eq. 1 (see Materials and Methods). The results of this analysis are shown in Figures 4A and B (see also Figure S2). Notably, we found that the data satisfactorily fitted the percolation function (R^2^ values ≥0.91–0.97) and with *β* percolation exponents ranging from 0.69 to 0.82 (Figure 4B and Figure S2C-F), in remarkable agreement with the universal critical exponents for the dimensionalities of *n*=4, across most of the ring, and *n*=5 near the periphery (0.65 and 0.83 for *n*=4 and *n*=5, respectively) [32]. These results also suggest that cell-density-dependent synchronicity derives from the existence of a critical time, *t*_*c*_, at which the population switches its behavior abruptly. The above results with mutant *C. violaceum* CV026 cells begged the question of whether the QS response of wt *C. violaceum* displays a similar “percolation-like behavior”. Gallardo *et al.* [28] used highly sensitive spectroscopic techniques to measure violacein production *in vivo* by wt *C. violaceum*. Figure 4C shows that their data can be readily fitted to the percolation function (R^2^ = 0.992, see also Table S1), albeit with a smaller *β* percolation exponent (0.46). Despite the notable differences between their experimental setup and ours, these results confirm our suspicion that the standard QS response in wt *C. violaceum* cells, possessing both the “QS listening” and “QS speaking” modules, displays an all-or-none character similar to that described here for the “QS-speaking” deficient strain CV026.

**Figure 4.**
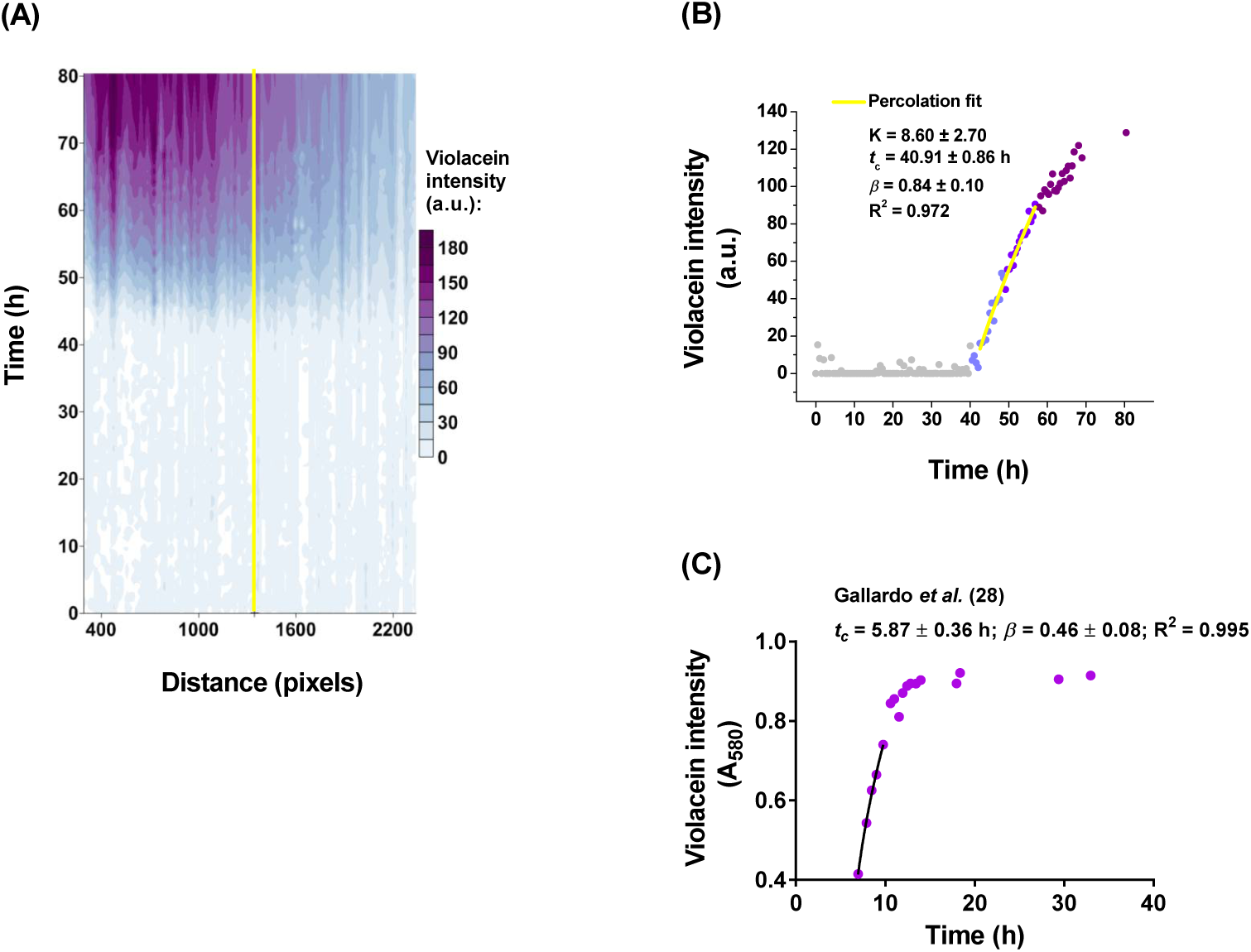
Adjustment of the density-normalized QS response of C. violaceum CV026 to the percolation function. A. Contour plot of violacein accumulation as a function of time and distance from the edge of the well. X-Axis: distance in pixels. Y-Axis: time. Violacein intensity in arbitrary units is represented according to the palette shown on the right. B. Fitting of the percolation function to the violacein accumulation data of panel A. The portion of the data used for the fit is indicated by the solid yellow line in A. The data corresponds to a single, representative experiment. C. Fitting of the data of Gallardo *et al.* [28] to the percolation function. Violacein intensity over time dispayed by wild type *C. violaceum*. X-Axis = time (h). Y-Axis = violacein intensity (A580). Note that the baseline for violacein intensity was set at A580 = 0.4 (see Materials and Methods and Table S1 for additional details).

The fact that our experimental data could be successfully fitted to percolation suggests that there is a correspondence between QS-regulated processes and PT. In this paper, we will make no attempt to rationalize the involved phenomena in terms of statistical mechanics, as has been done for other percolation phenomena. For the scope of this work, we will use percolation equations as a convenient mathematical tool to describe QS. For the sake of clarity, we will hereon use the terms “percolation phase” and “percolation kinetics” to refer to the behavior of the system in the vicinity of *t*_*c*_, and to its delayed nature, respectively.

### QS response of a synthetic E. coli biosensor

To better understand the kinetics of “QS listening”, we used a fluorescent *E. coli* biosensor carrying a synthetic genetic device based on the *luxR/luxI* QS genetic circuitry of *V. fischeri* [33]. In this biosensor, binding of LuxR dimers to the *luxI* pR promoter in the presence of externally added N-(3-oxohexanoyl)-L-homoserine lactone (3OC_6_HSL, AHL _*fisch*_) drives the expression of GFP [33]. Figure 5A shows the results of experiments performed with three different concentrations of externally added AHL _*fisch*_. The semi-logarithmic growth curves (inset in Figure 5A) show virtually no apparent growth differences due to AHL _*fisch*_ concentration and prove that the cultures remain in log phase for the duration of the experiment (notice that only incipient stationary phase was achieved at the late stages of growth). Notably, the density normalized fluorescence response, FI/OD600, increased with growth up to a maximum level whose magnitude is determined by the AHL _*fisch*_ concentration. Despite the fact that the lowest AHL _*fisch*_ concentration (5×10^−10^ M) was below the *k*_*Hill*_ (Figure S3), it was still possible to quantify the evolution of fluorescence. In general, increasing AHL _*fisch*_ concentrations resulted in steeper curves and greater maximum values of the density-normalized fluorescent response. The results of fitting these curves to the percolation function are shown in Figure 5B (R^2^ values ≥ 0.97; Table S2 and Figure S4). Remarkably, both *β* and *t*_*c*_ parameters assume values that lie within a very narrow range and remain essentially independent on the AHL _*fisch*_ concentration (Figure 5C and D, Table S2). Moreover, in the case of the *β* exponent the obtained values are very similar to those of the universal exponent of *n*-defined percolating lattices with *n*=5, 0.83 (Figure 5C) [34]. Figure 5D and Table S2 also show that for all AHL _*fisch*_ concentrations, the *t*_*c*_ value (*i.e.*, the critical time to trigger the response) is very short (*t*_*c*_= 7–10min). Since a GFP molecule can be translated in about 17 seconds during fast growth [35] and the estimated average time constant for GFP maturation is 6.3 min [33], the above *t*_*c*_ value strongly suggests that the system percolates immediately or very shortly after induction. These results are in agreement with those of Canton *et al.* [33], obtained with a synthetic *E. coli* biosensor carrying the same LuxR-driven GFP expression system used by us. When we fitted their results to percolation a *t*_*c*_ of 2.38 min was obtained (Figure 6A), in good agreement with our conclusion that the system enters the “percolation phase” immediately after induction.

**Figure 5.**
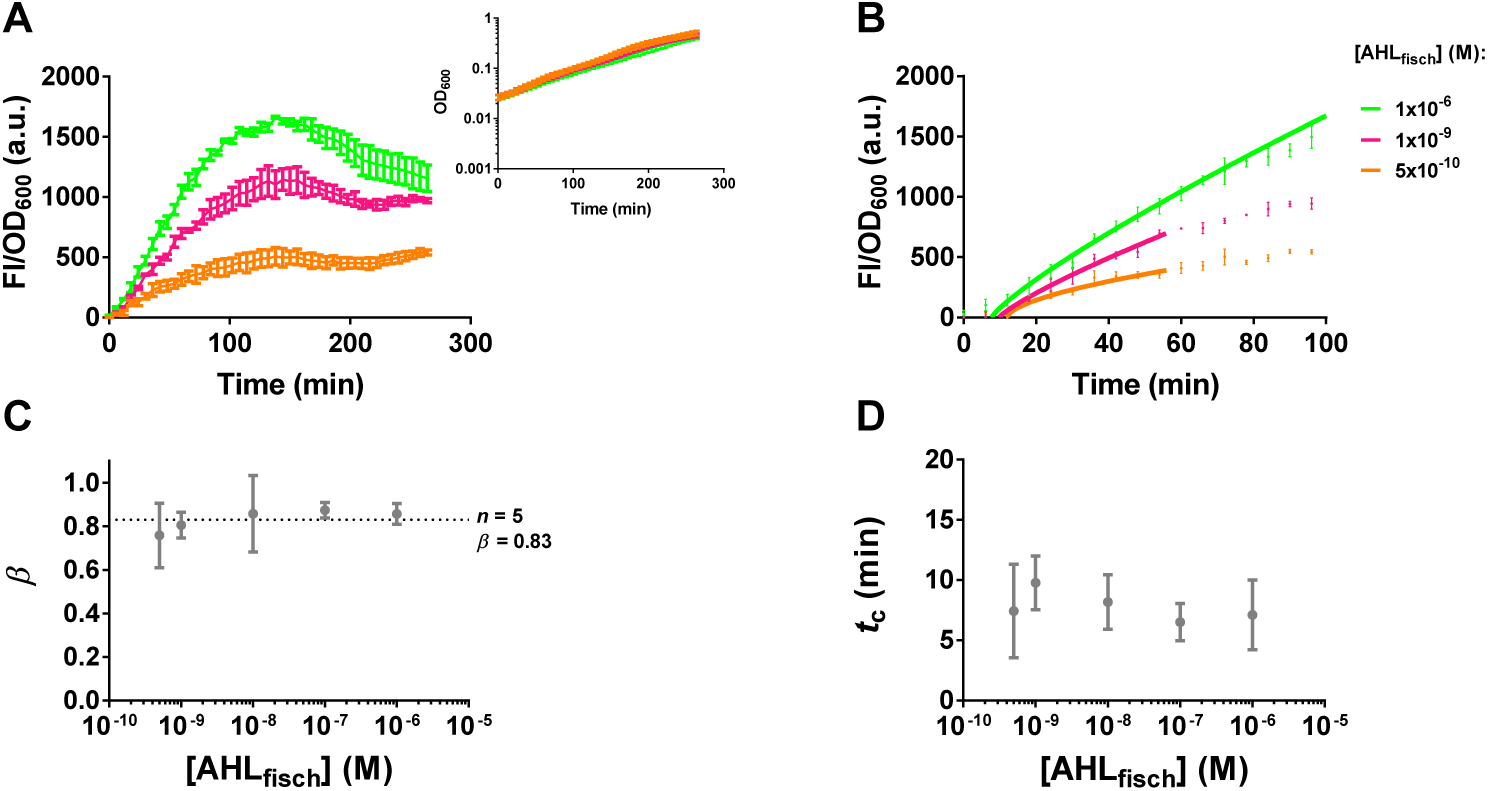
Adjustment of the density-normalized QS response of the E. coli biosensor to the percolation function. A. FI/ OD600 is shown as a function of time at three different AHL _*fisch*_ concentrations: 1×10^−6^ (green, above *k*_*Hill*_), 1×10^−9^ (magenta, close to *k*_*Hill*_) and 5×10^−10^ M (orange, below *k*_*Hill*_). Inset: Corresponding experimental OD600 at the three different AHL _*fisch*_ concentrations. B. Fit of the density-normalized QS response to the percolation function. The observed FI/OD600 values (solid circles) during the first 100 min of incubation at the three different AHL _*fisch*_ concentrations were fitted to the percolation function (solid lines). Panels C and D show the effect of AHL _*fisch*_ concentration on the percolation coefficients *β* and *t*_*c*_, respectively. The dotted line in panel C represents the value of the universal exponent for a fractal dimension value of 5 (*n*=5). Data represent the mean and standard deviation of three independent experiments with three biological replicates each.

**Figure 6.**
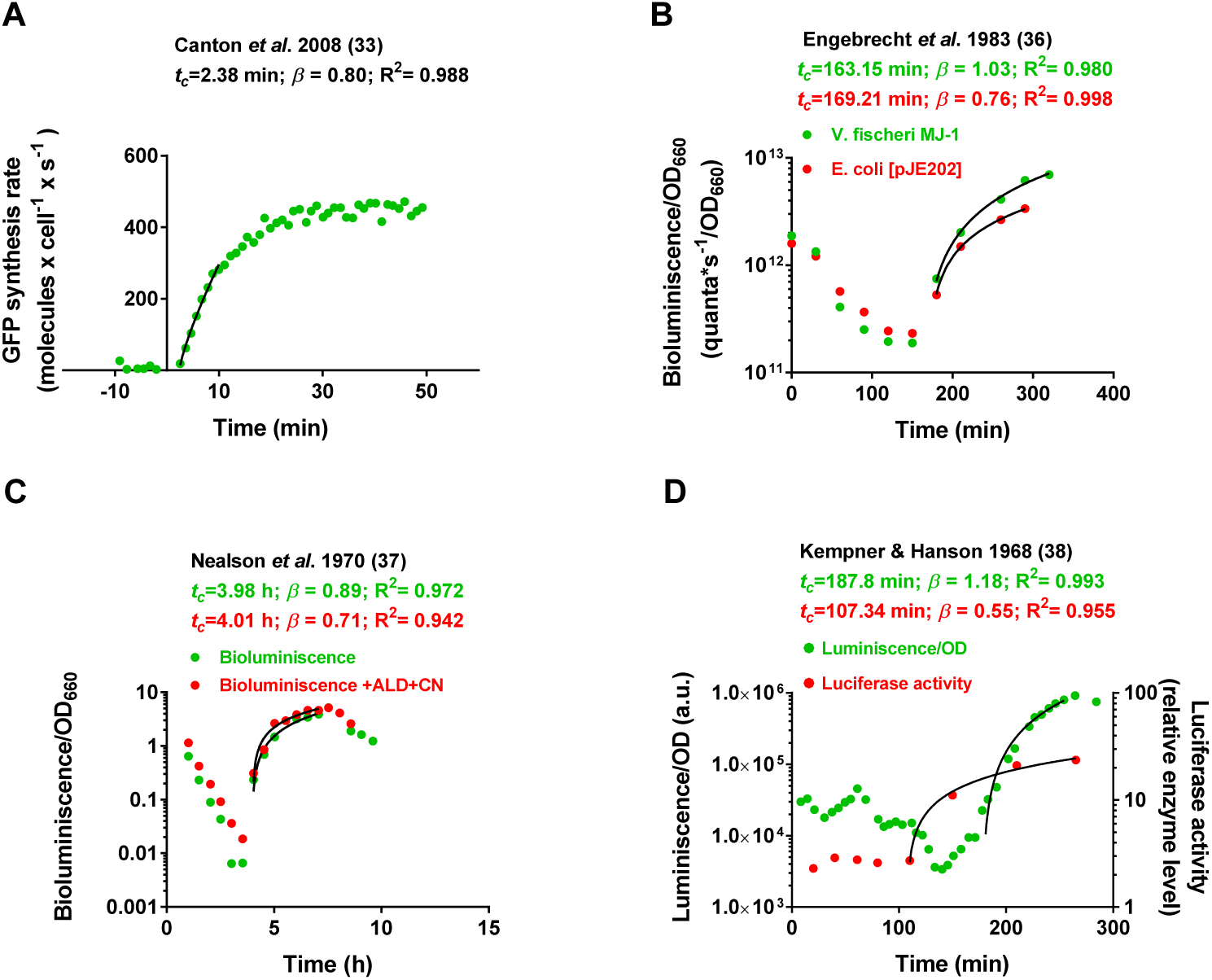
Adjustment of the density-normalized QS response of V. fischeri-based QS responses from the literature to the percolation function. A. Percolation fitting of the results of Canton *et al.* [33]. Cell-density normalized, QS-driven GFP fluorescence over time displayed by a synthetic *E. coli* biosensor. X-Axis = time (min). Y-Axis = GFP synthesis rate (molecules x cell^−1^ x s^−1^). Additional details in Table S1. B. Percolation fitting of the results of Engebrecht *et al.* [36]. Density-normalized luminescence yield over time displayed by *V. fischeri* MJ-1 (green circles) and *E. coli* pJE202 cells (red circles). X-Axis = time (min). Y-Axis = bioluminescence/ OD600 (quanta x s^−1^/OD660). Additional details in Table S1. C. Percolation fitting of the results of Nealson *et al.* [37]. Density-normalized, bioluminescence over time displayed by *V. fischeri* MAV cells. Experiments in the absence (green dots) and in the presence (red dots) of cyanide (CN) and aldehyde (ALD) are shown. X-Axis = time (h). Y-Axis = bioluminiscence/ OD660. Additional details in Table S1. D. Percolation fitting of the results of Kempner *et al.* [38]. Density-normalized luminiscence displayed by *V. fischeri* cells over time (green circles) and luciferase activity over time over time (red circles). X-Axis = time (min). Right Y-Axis = luminescence/OD (a.u.). Left Y-Axis = luciferase activity (relative enzyme level). Additional details in Table S1.

To glean an understanding of the reasons underlying the striking differences regarding the timing of the QS responses observed with the *C. violaceum* and the *E. coli* biosensors, we first searched the literature for examples of additional studies bearing relevance to our experimental setup. When the performance of the wt *luxR/luxI* device in its natural host (*V. fischeri*) is measured under experimental conditions that are comparable to ours, a delayed response ensued that could be fitted to the percolation function (Figure 6B, green trace, C and D; Table S1) [36–38]. Interestingly, a similar response has been measured in a recombinant *E. coli* strain carrying the wt *luxR/luxI* device on a medium copy-number plasmid (Figure 6B, red trace) [36] indicating that the immediate response of our *E. coli* biosensor is not related to the bacterial host itself. A major difference between the synthetic and natural devices is in the rate of expression of LuxR. In the case of our *E. coli* biosensor, the synthetic construct is carried on a high-copy-number plasmid, likely resulting in hundreds or thousands of copies of the luxR gene (http://parts.igem.org/Part:pSB1A3) [33]. In contrast, expression of *luxR* from its native promoter is expected to result in markedly lower levels of LuxR. While the actual intracellular levels of LuxR have not been measured, it has been estimated that it may range from ∼50 in the absence of AHL _*fisch*_ to ∼500 at high AHL _*fisch*_ concentrations, when LuxR is expressed from its native promoter and carried on a medium-copy-number plasmid [18]. Thus, the reason for the immediate response of our *E. coli* synthetic biosensor could stem from the existence of a saturating intracellular amount of QS activator at the outset of the assay, as opposed to cells with native QS genetic circuits, in which the levels of expression of these proteins would be below saturation.

We also imaged the QS response of our *E. coli* biosensor, as shown in Figures 7A–D. Strikingly, all the cells display QS-dependent fluorescence upon induction with AHL _*fisch*_ at concentrations below and above the calculated *k*_*Hill*_ (Figures 7A-D, Figure S3). While these are steady-state results and as such, do not reflect the synchronicity of the response, its simultaneous nature is captured in Movie S4. These results show that, in contrast to the stochasticity observed with *C. violaceum* CV026 at the single-cell level, our *E. coli* biosensor lacks variability in its response. Since high levels of single-cell variability has been described with “mute” *V. fischeri* MJ11 cells, carrying the native LuxR architecture [39], it follows that LuxR levels are also crucial in determining the stochasticity of the system.

**Figure 7.**
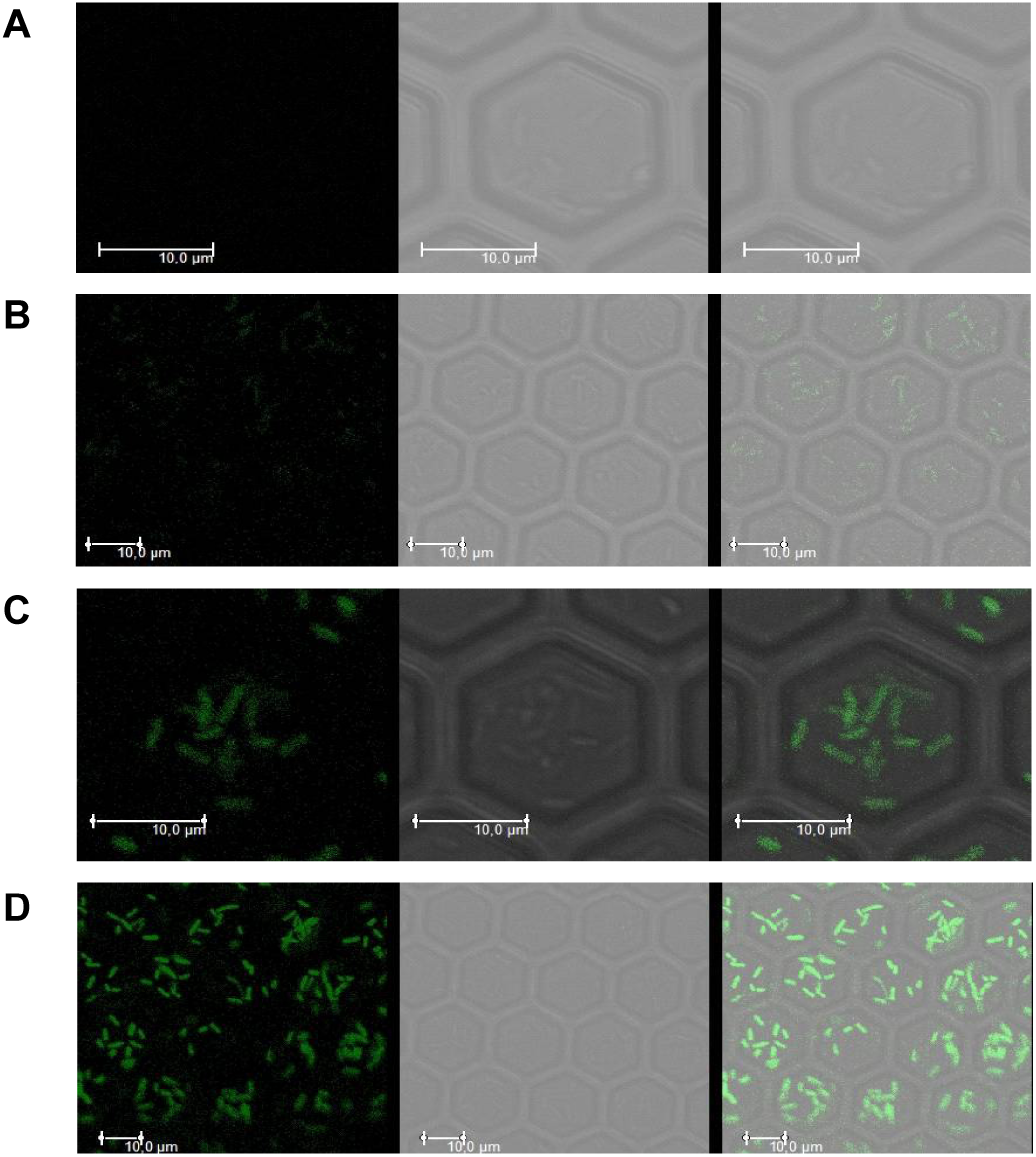
Fluorescence microscopy imaging of the E. coli biosensor. Cells were imaged in the presence of 0 (A), 2.5×10^−10^ (B), 1×10^−10^ (C), and 1×10^−6^ M AHL _*fisch*_ (D). The left image shows GFP fluorescence; middle image shows bright-field, and right image shows the overlay.

### The critical percolation time is sensitive to QS inhibitors

To test whether changes in the effective concentration of the QS activator could affect the timing of onset of the QS response, *t*_*c*_, we used the QS inhibitor *trans*-cinnamaldehyde (CA). CA is thought to compete with AHL and to block LuxR functionality by decreasing its DNA-binding ability [40, 41]. Figure 8A shows the results of a representative trial in which increasing concentrations of CA were added to the QS biosensor in the presence of a constant concentration of AHL _*fisch*_ (5×10^−10^ M). Notably, the all-or-none nature of the response, displaying a clear “percolation phase”, was maintained in the presence of the inhibitor. While we have observed variation from experiment to experiment (Figure S5), the general trend is highly consistent and shows a clear dose-dependent retardation of the response’s onset upon CA addition. Figures 8B and C show that *t*_*c*_ increases linearly with CA concentration, whereas the *β* exponent appears to be independent of CA concentration and affords values centered towards the universal value for *n*=5. Notably, CA can effectively delay the onset of the “percolation phase” in our QS system at concentrations that do not affect cell growth (< 250 µM CA) (Figure 8D). These results are consistent with the above idea that the intracellular levels of the QS activator follow an inverse relationship with *t*_*c*_.

**Figure 8.**
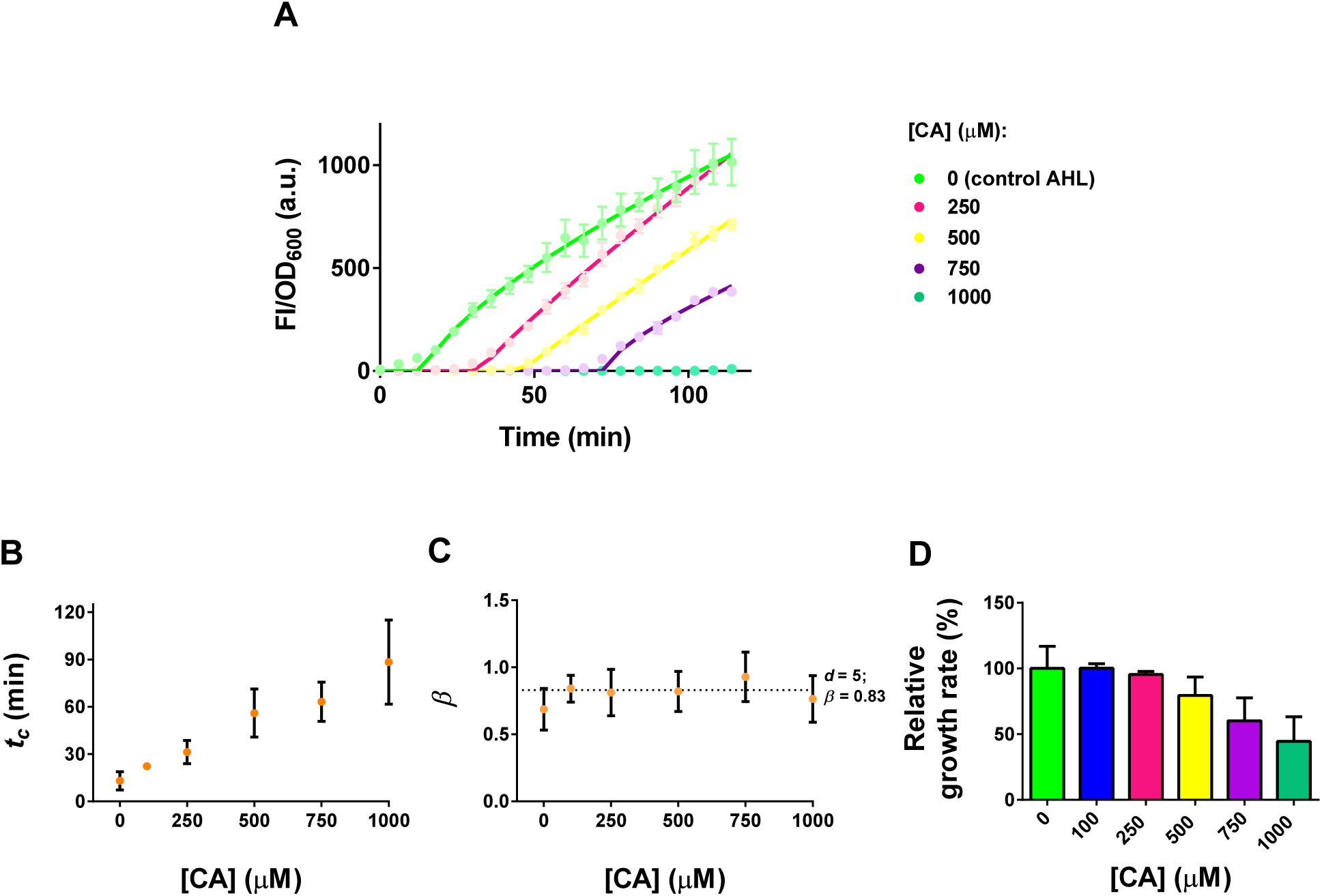
Effect of trans-cinnamaldehyde (CA) on the percolative response of the E. coli biosensor. A. Representative plot of FI/OD600 values (solid dots) as a function of time for the *E. coli* biosensor after treatment with varying CA concentrations in the presence of 5×10^−10^ M AHL _*fisch*_. Solid lines represent the best fit to the percolation function. CA concentration is shown to the right. Panels B and C show the effect over time of CA on the percolation parameters *t*_*c*_ and *β*, respectively. The plots were generated with the data shown in panel A and in Figure S5. The dotted line in panel C represents the value of the universal exponent for a fractal dimension value of 5 (*n*=5). D. Effect of CA concentration on the growth rate of the *E. coli* biosensor during the first 100 min of incubation. Data represent the mean and standard deviation of three to six independent experiments with three biological replicates each.

### *In silico* modeling of QS

To glean insight into how microscopic events at the single-cell level can lead to cell-density-dependent synchronicity at the macroscopic level, we generated an *in silico* model of “QS listening”. Since a comprehensive *in silico* model of QS would require us to measure all the relevant microscopic, biochemical parameters, a situation far from realistic in the case of *C. violaceum*, we decided to simplify our simulations by taking advantage of the fact that “QS listening” can be approximated by PT. The endpoint measures of success of our *in silico* model would be as follows. 1) The modeled response must fit a percolation function of the form of Eq. 1 (see Materials and Methods). 2) When running in “*C. violaceum* mode” the model must recreate the delayed all-or-none response observed with the *C. violaceum* CV026 biosensor. 3) The model must reproduce the stochasticity associated with violacein production by the *C. violaceum* CV026 biosensor. 4) Likewise, when running in “*E. coli* mode” the model must be able to reproduce the immediate response observed with the *E. coli* synthetic biosensor and its universal character at the single-cell level. 5) The model must be sensitive to the presence of antagonist inhibitors of QS. 6) In both running modes, the model must recreate the Hill behavior of the experimental data.

Following up on the results of Kurz *et al.* [18], we aimed at recreating a system of virtual cells in which the intracellular levels of the QS activator stochastically accumulated within each bacterium in the presence of AI. To do this, we conceived a simple, open-ended computational model of “QS listening” describing the growth of idealized bacterial cells in a 3D box (see Materials and Methods and Figure S6). Within this box, bacteria can interact with external AHL according to user-assigned parameters ascribed to the microscopic rules governing “QS listening”, *i.e.* growth rate, promoter copy number, probability of activator synthesis, AHL binding, activator dimerization, etc. The algorithm would then simulate the flow of molecules through the system, given a set of initial parameters. For the sake of simplicity, concepts such as transcriptional and translational bursts as well as the existence of a feedback loop at the level of the QS activator were intentionally excluded from the model.

First, we ran our simulations in “*E. coli* mode”, *i.e.* with hundreds of activator monomers at time zero and a relatively large monomer degradation constant in the absence of AHL (Table S3) [42]. We measured activator dimerization as a proxy for the ensuing QS response. Figure 9A shows that under these conditions, the presence of a large number of activator monomers at the time of induction results in a quick rise in the number of dimers and generates a response curve closely resembling that of the *E. coli* biosensor (Figure 5A). The fact that cell-density-normalized, *in silico* responses could be successfully fitted to the Hill function (compare Figure S7A to Figure S3), gave us additional confidence in the agreement between the *in silico* and the *in vivo* data. This was further corroborated by the successful fitting of the simulation data to a percolation function with a *β* exponent and a *t*_*c*_ that are in close agreement with the values measured experimentally (Figure 9A).

**Figure 9.**
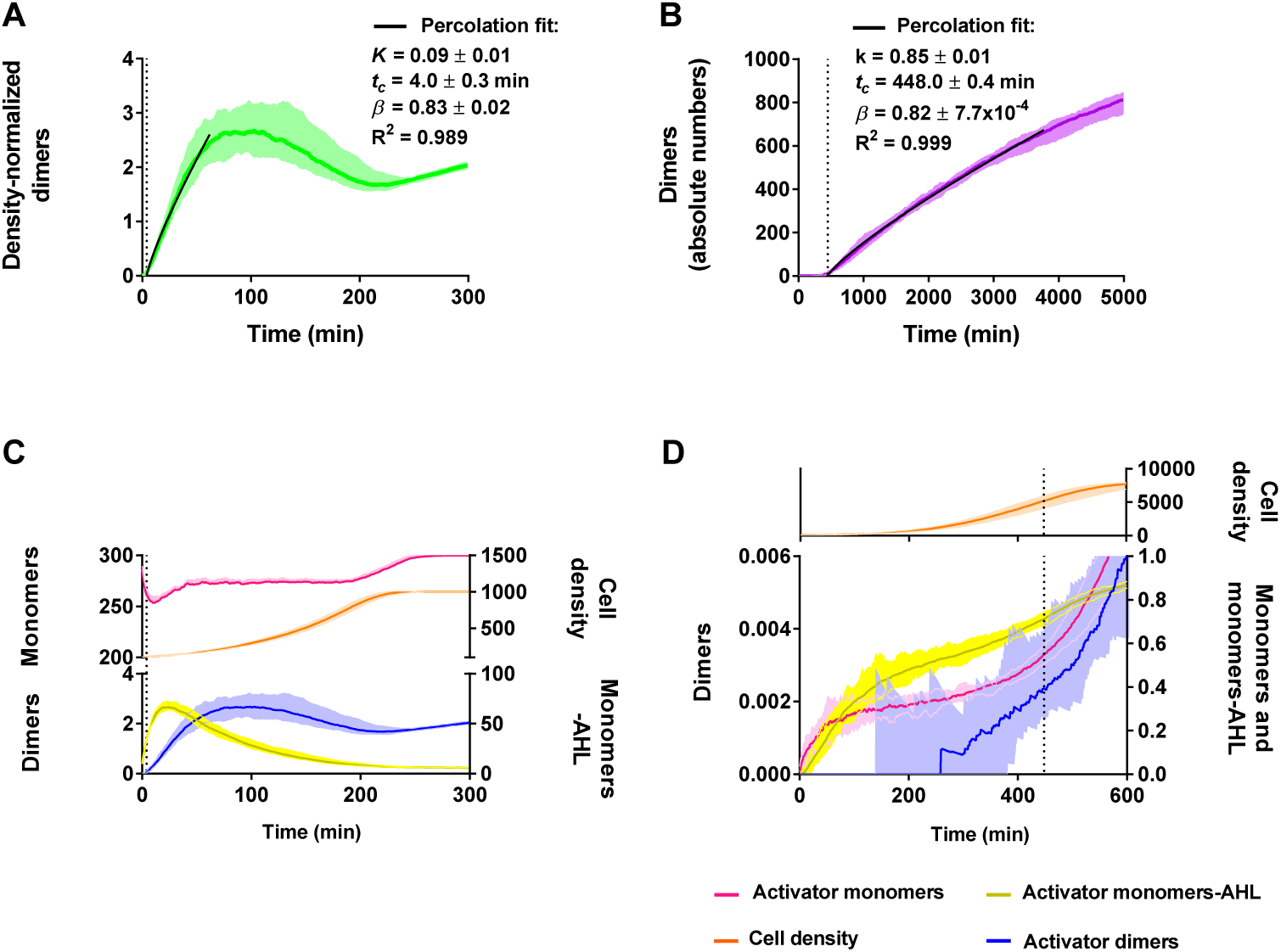
“QS listening” simulations in the “E. coli” and “C. violaceum modes”. A. Simulated accumulation of density-normalized LuxR dimers as a function of time in “*E. coli* mode”. X-axis: time in min. Y-axis: density-normalized number of LuxR dimers. The median of 10 simulations is represented by the solid green line and the data range is indicated by the light shadowed area. B. Simulated accumulation of CviR dimers as a function of time in “*C. violaceum* mode”. X-Axis: time in min. Y-Axis: absolute number of CviR dimers. The median of 23 simulations is represented by the solid purple line and the data range is indicated by the light shadowed area. The solid black line in A and B shows the best fit to the percolation function. C Density-normalized levels of activator monomers and dimers obtained in “*E. coli* mode”. Upper left y-axis: density-normalized monomer levels. Lower left y-axis: density-normalized dimer levels. Upper right y-axis: number of cells in the virtual box. Lower right y-axis: density-normalized monomer-AHL levels. D. Density-normalized levels of activator monomers and dimers obtained in “*C. violaceum* mode”. Lower left y-axis: density-normalized dimer levels. Upper right y-axis: number of cells in the virtual box. Lower right y-axis: density-normalized monomer and monomer-AHL levels. Data represents the median of 10 (C) and 23 (D) simulations and the data range is indicated by the light shadowed areas.

Recreating the response of *C. violaceum* CV026 was more challenging. Since each bacterium contains a single chromosomal copy of the gene encoding the QS activator CviR, it is likely that very few copies of CviR exist in *C. violaceum* CV026 in the absence of external AHL _*viol*_. This is because QS activators are unstable in the absence of their cognate AHLs [42]. Since it is known that macromolecules at single- or low-copy numbers behave stochastically in living cells [43], we wondered whether simulating the stochastic expression from the single *C. violaceum cviR* gene could give rise to a cell-density-dependent response, similar to the one observed *in vivo*. After testing different combinations of simulation parameters, we found that if activator levels were allowed to increase slowly in the cell (see simulation parameters in Table S3), a QS response with features similar to those measured *in vivo* was obtained (Figure 9B). Indeed, fitting of the simulation plots to the percolation function was successful, yielding a *β* exponent of ∼0.8 which is in good agreement with the experimental values obtained *in vivo* (Figure 4B). A *t*_*c*_ of 448 ±0.4 min places the onset of the QS response near the end of log-phase, in agreement with our own observations (not shown) and measured QS responses [44]. This is quite remarkable, as no growth-phase constraints were implemented in the model.

*In vivo*, the QS response of *C. violaceum* CV026 cells displayed a remarkable degree of sigmoidicity relative to the concentration of AHL _*viol*_ (Figure 1B). To test whether a similar behavior could be observed *in silico*, we run our simulations at different AHL concentrations. When the results are arranged in a Hill plot, a sigmoid curve showing a plateau that spans a wide range of AHL concentrations is apparent (Figure S7B). These data indicate that our algorithm can recreate conditions similar to those observed in plates of *C. violaceum* CV026.

When the results of both types of simulations are compared, a clear picture emerges. In agreement with our working hypotheses, both responses displayed “percolation kinetics” and their onset was consistent with the experimental evidence (“*E. coli* mode”: *t_c_* ∼4 min *vs.* “*C. violaceum* mode”: *t_c_* ∼7.5 h; Figures 9A and B, respectively). Attempts to achieve closer resemblance to the experimental results were not pursued at this stage of refinement of the algorithm. Notably, in both cases, the system enters the “percolation phase” once a small number of stable activator dimers emerge out of an intracellular monomer-AHL population that is much larger than that of dimers. Specifically, the levels of monomer-AHL complexes measured at *t*_*c*_ exceed by ∼2.5 and ∼1.5 orders of magnitude the amount of dimers in “*E. coli*” and “*C. violaceum* modes”, respectively (*c.f.* panels C and D in Figure 9).

In compliance with the above requirements, our *in silico* model was able to recreate many aspects of the QS response observed *in vivo*. For example, we attempted to model the effect of CA *in silico*, by assuming that the inhibitor worked as an antagonist of AHL [40]. Initial trials with the “*E. coli* mode” parameters failed due to the impossibility to simulate enough antagonist molecules to recreate the concentration range (relative to AHL) used *in vivo*. To solve this problem, it was necessary to sharply reduce the amounts of signal molecules, activator promoters, and of activator molecules present in the system at time zero (Table S3). Despite the changes, the overall shape of the response did not differ much from that obtained with the previous simulation parameters (*c.f.* Figure 9A to the curve with no antagonist of Figure 10A). Similar to the *in vivo* situation, we observed a clear retardation of the response’s onset whose magnitude correlates well with the concentration of the antagonist (*c.f.* Figure 10A to Figure 8A). This behavior was associated with a sharp decrease of intracellular activator-AHL monomers (Figure 10B) leading to the formation of fewer activator dimers as the antagonist concentration rose (Figure 10A and C). In agreement with our initial hypothesis on the role of LuxR-like activators, when the levels of activator-AHL monomers are plotted against *t*_*c*_, a clear reciprocal relationship between the two parameters is apparent (Figure 10D).

**Figure 10.**
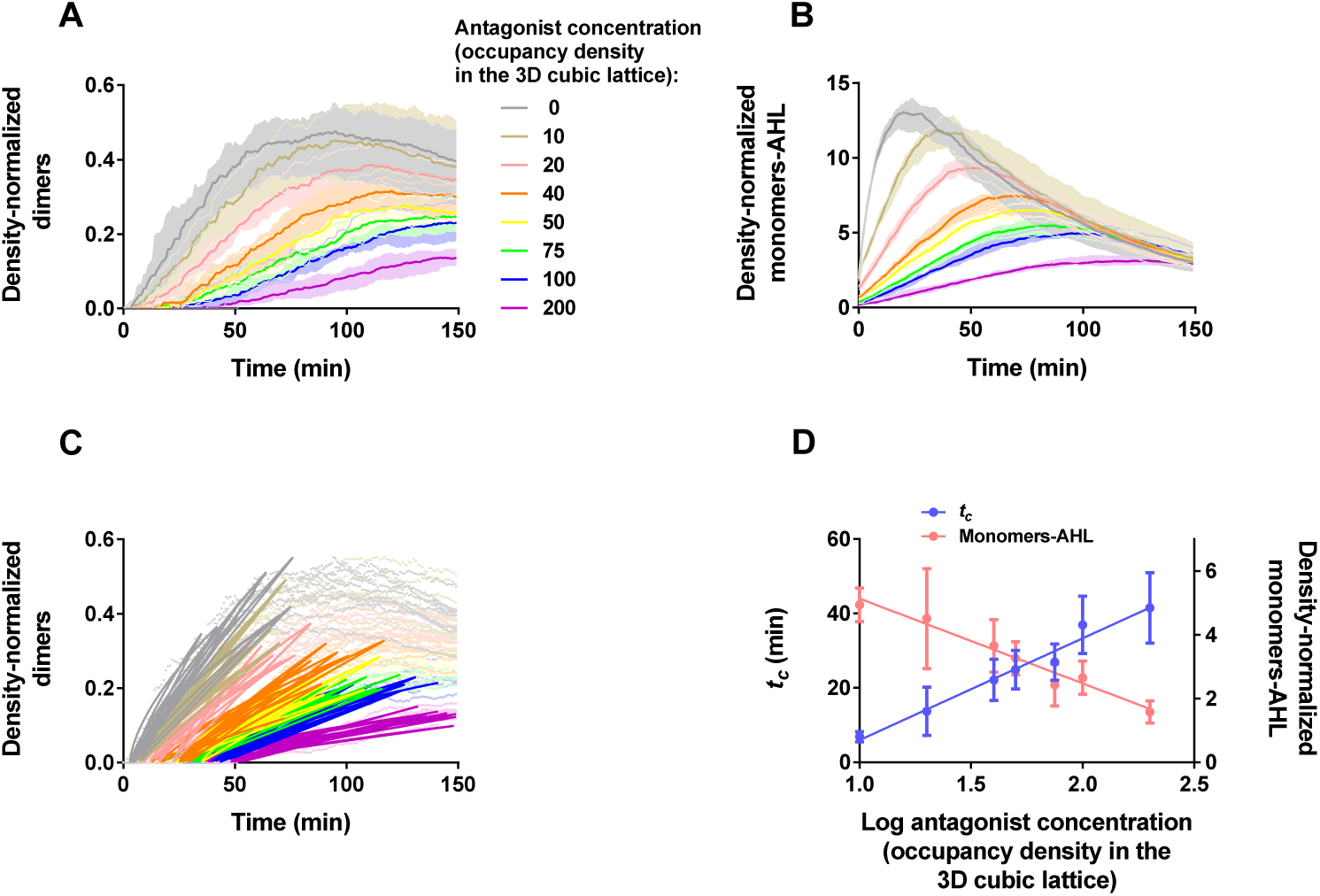
Simulation of QS in the presence of antagonist. A. Simulated accumulation of density-normalized LuxR dimers as a function of time in “*E. coli* mode”, in the presence of increasing concentrations of antagonist. The median curves of ten simulations have been color coded to the antagonist concentrations indicated to the right. The range is shown as a shaded area surrounding each curve. B. Cell-density normalized accumulation of monomer-AHL as a function of time in the presence of increasing concentrations of antagonist. Color coding and ranges as in A. C. Percolation fitting of activator dimer accumulation. Fitting traces indicated by solid lines, color coding as in A. D. Plot of *t*_*c*_ (light blue) and critical monomer-AHL concentration (salmon) *vs.* log of antagonist concentration.

The algorithm was also capable of pinpointing important microscopic differences between the modeled biosensors. Figures 11A and B, show that whereas activator dimer formation occurs in a minority of cells in the “*C. violaceum* mode”, virtually all the cells in the “*E. coli* mode” carry dimers at the end of the simulation. These differences in the number of cells implicated in the response explain why the density-normalized levels of QS activator species are much smaller in “*C. violaceum* mode” than in “*E. coli* mode” (*c.f.* panels D and C in Figure 9).

Since our simulations track the “activation” of individual virtual cells, the fact that their responses could be approximated by PT suggested that our percolation function can faithfully capture the integration of the stochastic “activation” times displayed by individual cells into a single population response. Weber and Buceta [19] have recently looked at the epigenetic variability of QS *in silico* by exhaustively tracking the individual trajectories of virtual single cells. Thus, their results offer an additional opportunity to test the above hypothesis regarding the connection between PT and QS. The fact that we were able to fit their population average response to PT (Figure 11C), clearly supports the existence of such connection. In other words, in the context of QS, PT connects single-cell stochasticity to the average population behavior.

**Figure 11.**
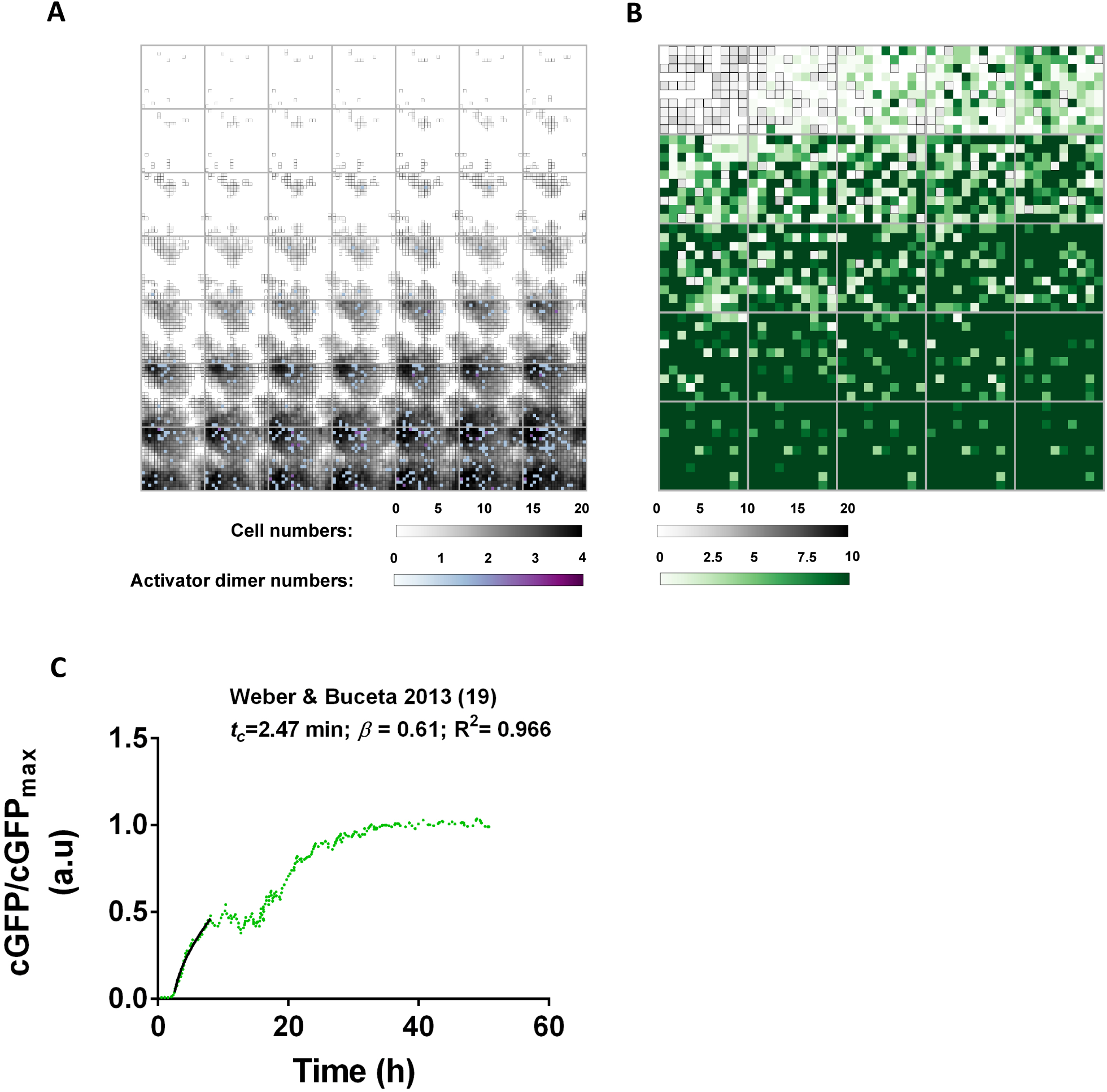
Population vs. single-cell responses. 2D representation of a simulation run in “*C. violaceum* mode”, showing the accumulation of activator dimers over time in the virtual bacterial population. B. 2D representation of a simulation run in “*E. coli* mode” showing the accumulation of activator dimers over time in the virtual bacterial population. Each tile corresponds to one of 49 (“*C. violaceum”* mode) or 25 (“*E. coli* mode”) time steps equally spaced from time zero until the end of the simulation. The plots are the result of the projection of the apical view of the 3D cubic lattice into 2D. Each pixel corresponds to a column in the box. Column cell density as a function of time is represented in gray scale. Column accumulation of LuxR dimers is represented in purple (A) and in green (B). The color gradients indicate the number of cells and dimers per column in the two simulation “modes”. C. Percolation analysis of Weber and Buceta’s data [19]. Accumulation of normalized, QS-driven GFP fluorescence over time in Weber and Buceta’s *in silico* model of [19]. X-Axis = time (h). Y-Axis = cGFP/cGFP*max* (a.u.). See additional details in Materials and Methods and in Table S1.

In summary, the fact that our open-ended computer model can recreate so many aspects of the experimentally observed QS responses attests to the robustness of our algorithm. At the same time, the success of our simulations reveals the simple nature of the microscopic rules governing “QS listening”.

## Discussion

Kempner and Hanson described in their seminal 1968’s paper an inverse relationship between the initial cell density of *V. fischeri* cultures and the delay of the observed QS response [38]. This and similar observations eventually led to the idea that the synchronicity of QS responses is dependent on cell-density sensing by individual cells. Another crucial piece of information in this puzzle was the finding that QS responses were always observed late during logarithmic growth [28, 30, 31, 38, 44]. To explain these phenomena, the classic and still current view of QS claims a master role for AI molecules, whose accumulation up to a threshold level determines the timing of the response [7]. In this paper, we have provided evidence that challenges this AI-centered view of QS and establishes LuxR-like activators as the master coordinators of bacterial cell-to-cell communication. Being QS the most basic form of cell-to-cell communication, these results have important implications for our understanding of the evolution of language in its primordial chemical form.

We have shown that the establishment of synchronized QS responses with similar temporal kinetics and cell-density dependence of standard QS responses do not require AI synthesis and as such, is independent of cell-to-cell communication. This implies that the ability to trigger such responses must be ingrained in the “QS listening” module. Clearly, such a realization has to be reconciled with models aimed at explaining QS. Furthermore, we have shown that, while highly coordinated in time, QS responses need not involve a large fraction of the bacterial population. Thus, a successful QS model must also account for the minoritarian character of the response observed at the microscopic level *vs.* the all-or-none nature observed macroscopically.

How can bacteria sense population levels in the absence of cell-to-cell communication? The classical AI-centered model of QS, claiming that AI accumulation up to a threshold level is the only determinant for the appearance of an all-or-none QS response, fails to explain our results. Such a system would require the presence of steady intracellular levels of activator molecules, passively waiting until the critical AI concentration is reached and the system can be driven into an all-or-none response. In light of the results presented here, there are several problems with this scenario. First, the presence of steady intracellular levels of activator molecules at time zero is completely inconsistent with the observed response of *C. violaceum* CV026. The formation of CviR dimers at time zero due to the presence of saturating concentrations of AHL _*viol*_ would result in an immediate response similar to that of the *E. coli* biosensor. Clearly, this is not the case, as the QS response of *C. violaceum* CV026 is delayed by many hours in a cell-density-dependent manner (Figures 2A and B). A second problem is related to stochasticity. Clearly, the presence of steady intracellular levels of activator molecules would eliminate stochasticity at the level of the QS activator. On the other hand, the presence of saturating levels of AHL _*viol*_ at time zero also eliminates any possibility of stochastic behavior due to potential differences in local AI levels. Thus, a new layer of regulation would have to be called into place to account for the stochastic character of the response observed at the microscopic level (Figure 3C). While this possibility exists, the well-known direct regulation of *vioABCD*, the operon of *C. violaceum*, responsible for violacein biosynthesis, by CviR [25], strongly argues for a simpler explanation. Taking all this into consideration, it stands to reason that an AI-centered model of QS fails to explain the cell-density-dependent response observed with *C. violaceum* model of “QS listening”.

Regarding stochasticity, both our *in vivo* and *in silico* results, together with those of others [18], are in agreement with the idea that the phenotypic or epigenetic variability is inversely related to the initial intracellular levels of QS activators (*c.f.* the responses of the *C. violaceum* CV026 and *E. coli* biosensors in Figures 3C and 7A-D, respectively). Two key experimental observations made with our *C. violaceum* CV026 biosensor provide essential clues to understanding how phenotypic variability can be achieved in the absence of cell-to-cell communication. The steady accumulation of intracellular activator-AHL molecules up to a threshold level, *ac*, predicted by our *in silico* model when running in “*C. violaceum* mode” (Figure 9D), could be interpreted as the action of a simple timer. However, the aforementioned inverse relationship between the initial cell density and the delay of the QS response observed *in vivo* with *C. violaceum* CV026 cells (Figure 2B) does not support this simple view. The fact that all cells come from the same original overnight culture and should have comparable intracellular distributions of CviR, completely rules out the timer hypothesis. Instead, our *in vivo* results are in agreement with the idea that the dilution of the intracellular pool of CviR-AHL _*viol*_ upon cell division effectively excludes the possibility of reaching the threshold level, *ac*, until a certain cell density has been reached. Decreased growth due to entry in stationary phase could easily explain this observation. By slowing down the dilution process, the intracellular levels of activator could quickly reach *ac* in the cell lineages carrying higher, stochastically driven concentrations of activator-AHL monomers. This, together with the fact that no violacein is produced by stationary-phase cells even in the presence of saturating amounts of AHL _*viol*_ (Figure 2C), also explains how epigenetic variability is achieved (Figure 3C). The well-known arrest of protein synthesis during stationary phase [45] should effectively prevent the accumulation of additional activator-AHL monomers in non-growing cells. Hence, cell lineages carrying lower levels of activator-AHL monomers will never reach *ac*. It should be noted that despite the fact that no stationary-phase specific rules were implemented in our *in silico* model, it successfully simulated a viable solution for the operational behavior of a natural “QS listening” module such as the one of *C. violaceum*.

By comparing the disparate responses of our *C. violaceum* and *E. coli* biosensors, together with the re-examination of relevant examples available in the literature, we have proposed that the intracellular levels of LuxR-like activators are responsible for all the observed differences. The overall agreement between our *in silico* and *in vivo* results strongly suggests that the basic principles and the logic behind our modeling strategy are likely correct. Bearing this in mind, we may take our *in silico* results as initial constraints for a “low-resolution” model of the molecular interactions and the epigenetic events governing the buildup of the “QS ear”. The model is schematically depicted in Figure 12. The generation-1 bacterium shown in Figure 12A stochastically synthetizes a LuxR-like monomer (blue ovals) following the firing of the stochastically-driven promoter, *P_SD_*, controlling its expression (leftward, black arrow in the magnified view of the generation-1 bacterium in B). Following cell division, the monomer is inherited by one of the two daughter cells (generation 2). The presence of saturating levels of external AI will ensure that the activator will be present in its monomer-AI form (AI, yellow pentagons in Figure 12A). The maintenance of stochastic expression over several generations (denoted by dots in Figure 12A), leads to varying levels of activator monomers in different branches of the lineage tree. This is similar to the concept of epigenetic memory proposed by Kurz *et al.* [18]. Due to the slowing down of growth at the end of log phase, generation *j* cells have more time to accumulate new activator monomers. As a result, the monomer-AI pool within the most monomer-enriched lineages will reach the activator threshold, *ac* and the cells will become QS-activated (purple cell on the bottom branch of the lineage tree). Upon reaching *a_c_*, the binding of an activator dimer (green ovals) to the rightward, QS-regulated promoter, *PSD*, allows the expression of QS-regulated genes (see magnified view in Figure 12C). Due to the slower growth rate, the activated state might be maintained in the progeny of this bacterium in other bacterial lineages that became activated during generation *j*. At the population level, a highly synchronized, cell-density dependent QS response with “percolation kinetics” becomes evident shortly afterwards. This is represented by the percolation curve starting in generation *j* and extending to generation *n* in A. Regarding monomer-poor lineages, translation arrest during stationary phase will ensure that these lineages will not become QS-activated even after prolonged incubation (top branch in Figure 12A). Viewed in this light, our *in silico* simulations, showing very small dimer/monomer-AHL ratios at the onset of the QS response (Figures 9C and D), lead us to propose that dimerization, and the ensuing QS response, requires the buildup of a rather high concentration of activator-AHL molecules. In other words, following a steady accumulation of activator-AHL monomers over several generations, a critical threshold level *a_c_* is reached upon which dimer formation ensues (Figure 9B and D). This result is in perfect agreement and indeed it explains the recent observation of Wang *et al.* [46], who showed that the highest sensitivity of LuxR-like activators is achieved at high intracellular concentrations of these molecules. The achievement of this threshold level coincides with the percolation critical time *t*_*c*_. In other words, the length of time required for the development of a synchronous QS response with “percolation kinetics” depends solely on the time necessary to build the “QS ear”, namely, an array of activator monomers large enough to drive dimerization in the presence of AHL. In the synthetic *E. coli* biosensor, the “QS ear” is immediately formed in all the cells, provided that enough external AHL _*fisch*_ is available. In *C. violaceum* CV026, however, the “QS ear” is built up over several generations. Despite the steady buildup, activator levels cannot achieve their threshold level, *a_c_*, until decreased cell growth overcomes their dilution rate. We expect that this “low-resolution” model will guide future research efforts aimed at elucidating the molecular details of “QS listening”.

**Figure 12.**
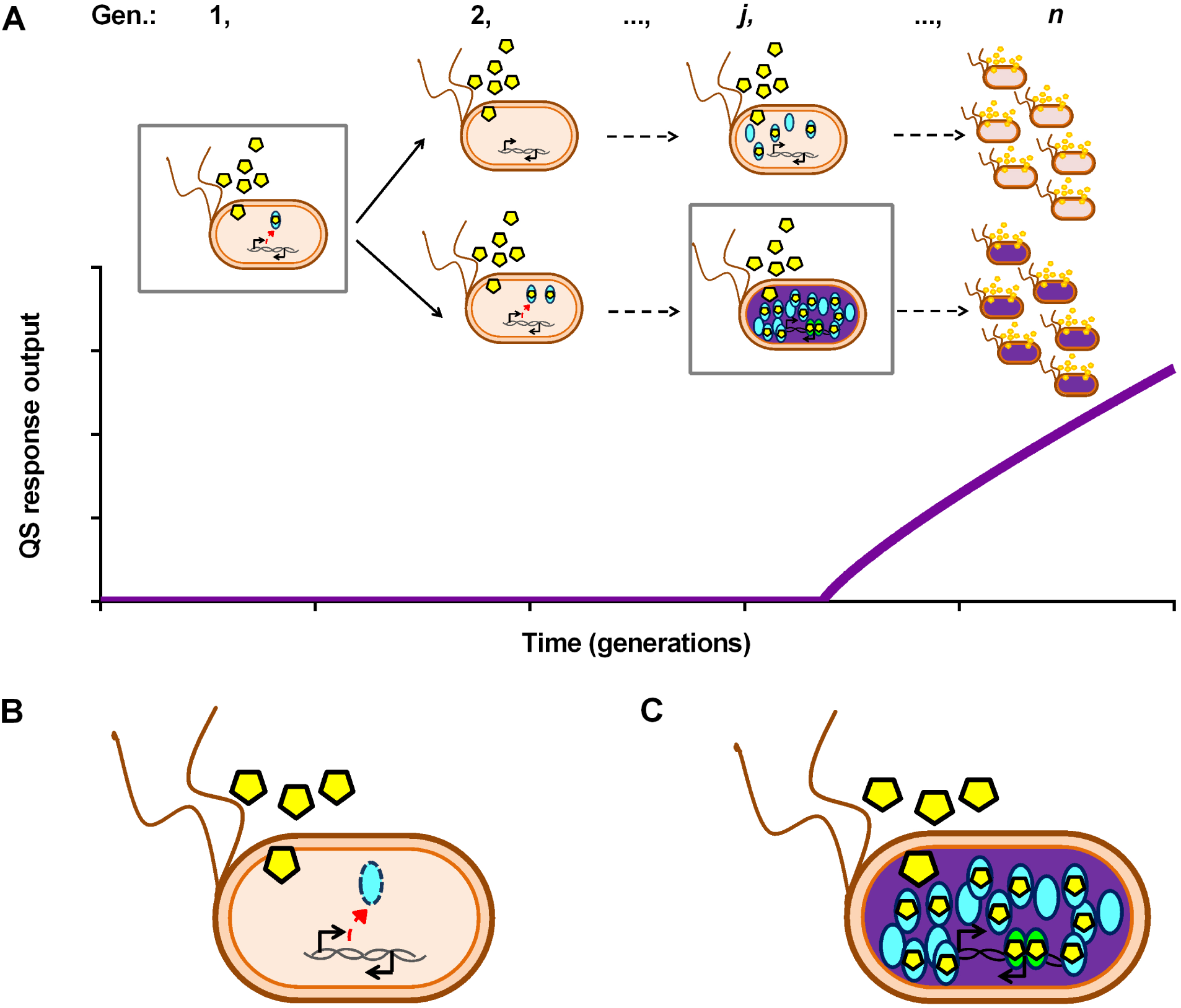
Schematic model of “QS ear” buildup. See main text for a detailed explanation. Bacteria are schematically shown as short rods with round ends. OFF bacteria are shown in pink, whereas ON bacteria are shown in purple. AHL molecules are depicted as yellow pentagons and activator monomers as cyan ovals. Bound AHLs are shown within activator ovals. Activator dimers are shown in green. Chromosomal DNA is depicted as a schematic double helix and the position of the two promoters, arranged in the *C. violaceum* orientation [47], is shown. The left promoter stochastically drives the expression of activator molecules, whereas the right promoter drives the expression of the QS response upon activator dimer binding. A. The fate of two cell lineages is shown over several generations. The different generations 1, 2…,*j*…,and *n* are indicated. A schematic percolation plot is shown under the cells to indicate the percolative character of QS. B. Detail of the boxed cells shown in A.

Before our studies, evidence obtained from recent *in silico* studies had already started to erode the tenets of the classical AI-centered model of QS. For example, Weber and Buceta [19] proposed that the molecular noise at the level of the QS activator, but not at the level of the AHL synthase, is what controls the precision of the QS switch. Their *in silico* experiments showed that fluctuations in the levels of the QS activator were responsible for the noise-induced stabilization of particular phenotypic states during the QS response. In agreement with this idea are the results of Kurz *et al.* [18], who demonstrated that epigenetic memory can be created by the noise-driven intracellular accumulation of QS activator molecules. Besides, these authors showed *in silico* that the presence of increasing concentrations of intracellular LuxR elicited a concomitant decrease in the delay of the responses’ onset (equivalent to our *t*_*c*_) [18]. These results are in perfect agreement with our view of LuxR-like proteins as the centerpiece of the QS response. In addition to this *in silico* data, some *in vivo* evidence existed that pointed towards an unsuspected role for QS activators. For example, the results of Williams *et al.* [13] showing that increasing the strength of the activator promoter *in vivo* tends to saturate the QS system with undesired copies of activator molecules, is in good agreement with our views. Similarly, Haseltine and Arnold [48] found that the timing of the QS response in an *E. coli* system with luxR expression originating from a low-copy-number plasmid (∼20 copies/cell) carrying the native *lux* architecture could be increased by decreasing the rate of luxR translation. In addition to these reports pointing out the central role of QS activators, other authors have directly questioned the importance of external AI levels. The single-cell studies of Pérez and Hagen [39], showing high variability in the QS response of “mute” *V. fischeri* MJ11 cells clearly contradict the concept that AI levels convey much useful information on the cell’s microenvironment. Similarly, Williams *et al.*’s results on QS hysteresis, *i.e.* the history-dependent behavior of QS systems, led to the proposal that endogenous synthesis of AHL would not be required for the presence of hysteresis per se, but would only affect the range of activator concentration at which hysteresis occurs [13]. Thus, it appears that two key features of QS, cell-density-dependent synchronicity and hysteresis do not rely on the ability of individual cells to sense population levels.

Being QS a type of critical phenomenon, it should not come as a total surprise that PT can be used to model its functioning. Recently, the Ising model for phase transitions has been proposed as a tool to explain cellular behavior arising from the workings of QS circuits [49]. These results, together with ours, prompt further investigation into the physical principles responsible for the mathematical description of the process of cell-to-cell communication as a phase-transition event. Our observations are indeed consistent with the notion that PT offers a simple and novel framework for modeling and mathematically interpreting the QS response. In favor of this hypothesis, we argue the following. First, the QS response has all the hallmarks of a critical “all-or-none” process when it is measured at the macroscopic level. Second, the calculated exponents after fitting experimental and simulated data to the PT function tend to fall rather close to conserved values corresponding to those of universal processes governed by percolation. Third, the “percolation phase” of QS responses is consistent with the integration of a multitude of noisy QS responses, observed at the individual-cell level, into a single population response measured macroscopically. Fourth, the ability of QS antagonists to delay the “percolation phase” in both our *in silico* and *in vivo* models of QS, strongly resembles the effect of inhibitors known to affect network formation in well-established percolation systems. For example, the rennet-induced coagulation of milk is driven by the enzymatic destabilization of the casein micelle and the subsequent flocculation and gelation of casein [50]. This phenomenon has been explained as a percolation phenomenon, whose *t*_*c*_ is inversely dependent on the concentration of the enzyme chymosin [51]. It has also been shown that gelation can be inhibited either by the addition of anions or by pre-crosslinking casein with transglutaminase [52]. Notably, in both cases, inhibition is associated to increased *t*_*c*_s, much like the QS behavior displayed by our *E. coli* biosensor in the presence of CA (Figure 8A).

A crucial aspect of PT is the issue of connectivity. At the present time, we lack evidence that can be used to establish how connectivity might be generated during QS. In the absence of such direct evidence, we can only speculate that connectivity could be started at the molecular level by the stochastic generation of high intracellular levels of activator-AHL monomers in some cells. The clonal propagation of these high levels of activator-AHL monomers during cell division could effectively result in the establishment of an interconnectable or percolable network of cells across the resulting cell lineages. In this regard, the expression rate of the QS activator and the cell’s growth rate would be the key control parameters dictating the probability of establishing connectivity in the system. Further research will be necessary to satisfactorily rationalize our observations in terms of statistical physics.

The fact that PT can be used to explain the development of QS responses deserves additional attention. Interestingly, two different quorum-related biological phenomena have been recently described in terms of PT. On the one hand, in the formation of neural synapses, stimuli from connected neurons are integrated into the target neuron, which fires once a quorum of stimuli is reached, in a way termed “quorum percolation” [22]. On the other hand, pheromone-regulated labor division in honeybees and other social insects has been compared with bacterial QS [53] and has been explained by PT [23]. Together with our results, these observations suggest that phenomena displaying “percolation kinetics” constitute the basis for the development of synchronized responses among cell collectives since the beginning of life.

The evidence presented here opens up the intriguing possibility that the “listening” skills could be acquired by bacteria independently of the ability to “speak” [6, 54–56]. For example, at a certain point in evolution, LuxR-like molecules could have developed the capability to sense quorum by binding to certain metabolites that naturally accumulated during cell growth [4]. The perfection of the mechanism by the incorporation of LuxI-like AHL synthases could be achieved in a subsequent evolutionary step. Such ideas would lead to the hypothesis that the most primordial form of language, could have evolved from the action of a single protein that was able to “listen in” on chemical cues present in the extracellular environment. Along these lines, the discovery of “solo” luxR elements, which are present in the genomes of some bacterial species in the absence of the usual QS circuitry [56–58], is all the more intriguing. The possibility that some of these elements might be remnants of primitive cellular functions from which the “QS ear” evolved should be considered. Alternatively, could these LuxR-like “solo” proteins be used by current bacteria to listen in on other cells by binding to some yet uncharacterized metabolite? [56]

In summary, our results lead us to propose that stochastic expression of QS activators can trigger delayed responses with cell-density-dependent synchronicity in the absence of AHL production. That nature has been able to exploit such a simple, noise-driven, genetic device for the development of coordinated responses in bacterial populations is indeed an astonishing evolutionary feat.

## Materials and Methods

### End-point evaluation of violacein production at various AHL concentrations

The mini-Tn5 mutant bioreporter strain of *C. violaceum CV026* has been described elsewhere [24]. Bioassay plates were made as follows. An overnight culture of CV026 was subjected to a 10^−1^ dilution in LB (final OD_600_ ≈ 0.4) and further sub-diluted 10^−1^ in a final volume of 3mL of semi-solid LB containing 1.2% agar and kanamycin (30µg/mL). The mixture was poured onto a LB agar plate supplemented with kanamycin (30 µg/mL). N-hexanoyl-DL-homoserine-lactone (Sigma-Aldrich; from now on referred to as AHL _*viol*_) was diluted to 1×10^−1^ M in acetonitrile, and the stock was stored at −20*°*C. Working solutions with concentrations ranging from 1×10^−2^ to 1×10^−6^ M were prepared by further dilution in water. Wells were formed into the agar at adequate distances, by using the wider end of a 1-mL micropipette tip. The wells were filled with 50 µL AHL _*viol*_ at concentrations ranging from 1×10^−5^ to 1×10^−6^ M. After incubation at 30*°*C for 48 h, high-resolution pictures were taken with a Fujifilm FinePix S3Pro camera (picture size= 1440 × 960 pixels, focal length= 70 mm). The pictures were analyzed with ImageJ 1.47v [59]. After splitting the image into RGB channels, the variation of color intensity in every violacein ring was measured on the green channel. To do this, six radial lines, each spanning 120–160 pixels were drawn starting at the edge of the AHL _*viol*_ source and extending across the imaged violacein ring and into the violacein-free periphery. Intensity values were corrected by subtracting the average pixel intensity of three lines (131 pixels each) drawn at the violacein-free periphery.

### Estimation of AHL diffusion profile from the calibration with bromophenol blue (BMB)

BMB was dissolved in water to a concentration of 0.9mM and filter sterilized. Three cell-free, bioassay plates were prepared as described above. A 50-µL aliquot of 0.9mM BMB was applied to the well, and the plates were incubated upright for 48 h at room temperature. BMB diffusion was recorded over 48 h, and the images were analyzed with ImageJ 1.47v [59]. To convert pixel intensity values to BMB concentration, a calibration curve was built by diluting BMB in semi-solid LB containing 1.2% agar to final concentrations ranging from 0.025 to 0.5 mM. The measured intensity values were fitted to a calibration curve of the form y = a – b*ln(x+c) (nonlinear curve fit Log3P1; OriginLab, Northampton, MA) (Figure S1A). The mixtures were overlaid onto LB agar plates and BMB intensity was measured in the RGB green channel as explained previously. The concentration of diffused AHL _*viol*_ in agar was estimated as follows. First, the concentration of BMB in the colored ring after 48 h diffusion in agar, plotted as a function of the distance from the edge of the well (Figure S1B), was fitted to an exponential function (OriginLab, Northampton, MA) of the form Y = Y0 + Ae^*R*o*X*^, (Figure S1C) thus yielding, Y = 9.29×10^−5^ + 0.13715 e^−0.01462*X*^, (R^2^= 0.991) where, 9.29×10^−5^ coefficient represents the best-fit estimated BMB concentration (in M) at the periphery of the ring after 48 h; 0.13715 is the estimated BMB concentration (in M) at the edge of the well (*i.e.*, at pixel position 0) after 48 h; and X is the distance in pixels from the edge of the well to the periphery.

### Time-lapse analysis of violacein production

For time-lapse imaging of violacein production in *C. violaceum* CV026, a 10^−2^ dilution of a *C. violaceum* CV026 overnight was plated onto bioassay plates as described above but with 50 µL of an aqueous 1×10^−6^ M solution of AHL _*viol*_ placed in the well. The evolution of violacein production over time was recorded at room temperature under a Leica S6D Greenough stereo microscope at the 6.3× zoom setting. The experiment was repeated two additional times at 30*°*C. To quantitate violacein production, a single radial line, spanning 2341 pixels, was drawn at each time step. Pixel intensities along the lines were plotted as a function of the distance, also measured in pixels (Figures 2A, 4A and S2). The intensity values at each time step were corrected by subtracting the pixel intensity of the line drawn at time zero.

### Time-lapse analysis of violacein production relative to cell density

Plates supplemented with AHL _*viol*_ were prepared as follows. Three mL of semisolid LB-agar (containing 17.5 g of agar per liter) with kanamycin (30 µg/mL) carrying 50 µL of 1×10^−2^ M AHL _*viol*_ were overlaid onto LB plates supplemented with kanamycin (30 µg/mL). Stationary phase overnights of *C. violaceum* CV026 were serially diluted (10^0^, 10^−1^, 10^−2^ and 10^−3^ dilutions) in fresh LB and 2-µL drops were placed onto the agar. Photos were taken at 30 min intervals (Figure 2B and Movie S2).

For the assays in liquid, 50 µL of 1×10^−4^ M AHL _*viol*_ were added to a 1.5-mL, stationary-phase culture of *C. violaceum* CV026 and to a 10^−2^ dilution of the same culture in fresh LB and incubated at 30*◦*C for 48 h (Figure 2C). No AHL _*viol*_ controls were incubated alongside.

### *E. coli* biosensor assays

The *E. coli* strain Top10 was transformed with plasmid pSB1A3 - BBa_T9002, carrying the BBa_T9002 genetic device (Registry of Standard Biological Parts: http://parts.igem.org/Part:BBa_T9002), kindly donated by Prof. Anderson Lab (UC Berkeley, USA). The transformed strain is a biosensor that can respond to the N-(3-oxohexanoyl)-L-homoserine lactone (named AHL _*fisch*_ hereafter).

To minimize experimental variability, we strictly adhered to the following seeding protocol. A flask with 10mL of Luria Bertani (LB) broth, supplemented with 200 µg/mL ampicillin, was inoculated with a single colony from a freshly streaked plate of strain Top10 pSB1A3 - BBa_T9002. After incubation for 18 h at 37*°*C with vigorous shaking, 0.5 mL aliquots of the overnight culture were mixed with 0.5 mL of 30% glycerol and stored at −80*°*C until further usage. Prior to each experiment, a glycerol stock from the single colony-culture was diluted by a factor of 10^−3^ into 20 mL of M9 minimal medium supplemented with 0.5% casamino acids, 1 mM thiamine hydrochloride, 0.4% glucose and ampicillin (200 µg/mL) and grown to an OD600 of 0.04 ± 7.76×10^−3^ (∼4 h). AHL _*fisch*_ (Sigma-Aldrich) was dissolved in acetonitrile to a stock concentration of 1×10^−1^ M and stored at −20*°*C until further usage. Prior to each experiment, AHL _*fisch*_ stock was serially diluted in water to yield working solutions ranging from 1×10^−2^ to 1×10^−9^ M. 20 µL of the AHL _*fisch*_ working solutions were transferred to the wells of a flat-bottomed 96-well plate (Greiner Bio-One, cat. #M3061). After the addition of 180 µL aliquots of the bacterial cultures, the final AHL _*fisch*_ concentration in the plates ranged from 1×10^−10^ to 1×10^−6^ M. Three blank wells with 200 µL of medium were used to measure the absorbance background. Additionally, three control wells were prepared to measure the fluorescence background. The plate was incubated in a Safire Tecan-F129013 Microplate Reader (Tecan, Crailsheim, Germany) at 37*°*C and with vigorous orbital shaking during the five seconds prior to each measurement. Absorbance and fluorescence were measured every 6 min with the following parameters: (fluorescence *λ_ex_*= 485 nm, fluorescence *λ_em_*= 520 nm, integration time= 40 µs, number of flashes= 10, gain= 100, measurement mode= top). To avoid complications due to excessive evaporation during the determinations, only 294 min of growth were recorded (measured evaporation = 6.7 ±0.27%; n=3). For each experiment, the fluorescence intensity (FI) and OD600 data were corrected by subtracting the values of absorbance and fluorescence backgrounds and expressed as the average for each condition (n=3). To estimate *k*_*Hill*_, the rate of evolution of FI/OD600 during the first 100 min of incubation was estimated from the value of the slope of a linear fit of FI/OD_600_. The dependence of the FI/OD_600_ rate as a function of AHL _*fisch*_ concentration was fitted to a non-linear growth sigmoidal Hill function by a minimization iteration process (OriginLab, Northampton, MA) and the best-fit parameters were determined (Figure S3). All measurements consisted in a minimum of three biological replicates.

### Microscopy

Low magnification images of *C. violaceum* CV026 plates such as those shown in Figure 3A and B were obtained by using the 40× zoom setting of a Leica S6D Greenough stereo microscope with an integrated photo port.

High magnification, bright-field imaging of *C. violaceum* CV026 cells (Figure 3C) was performed as follows. An overnight culture of *C. violaceum* CV026 was diluted by a factor of 10^−2^ into 5 mL of LB supplemented with kanamycin and 100 µL of 10µM AHL _*viol*_ and grown for two days at 26*°*C. Under these conditions, most of the dark purple color was found within a mucoid precipitate at the bottom of the tube. A loopful of this material was placed on a coverslip and imaged on a Nikon microscope with Plan Fluor 100X/1.30 DIC H/N2 ∞/0.17 WD 0.16.

Fluorescent imaging of the *E. coli* biosensor (Figure 7) was performed as follows. Cultures of the *E. coli* biosensor induced with various concentrations of AHL _*fisch*_ were prepared as described above. After incubation at 37*°*C with shaking for 300 min in the microplate reader, as described above, 200-µL aliquots were transferred to CytoCapture imaging dishes with 20-µm hexagonal cavities (Zell-Kontakt GmbH, Nörten-Hardenberg, Germany), followed by 30-min incubation at 37*°*C to allow individual cells to settle within the cavities. Confocal laser scanning microscopy (CLSM) was performed using a Leica TCS SP2 spectral confocal scanner mounted on a Leica DM IRES inverted microscope. All confocal images were acquired with the following settings: an HCX PL APO 63.0× 1.20 W CORR UV water-immersion objective, an argon excitation laser (488 nm), a 134.1 µM pinhole (1.0 Airy unit), standard Leica settings for GFP beam path, an emission bandwidth set to 500–600 nm, and a voxel width and height of 51.0 nm. Images were acquired from single scans with a line average of 4.0, scan speed of 400 Hz, and 8-bit resolution. Image processing was performed with Leica LAS AF Lite software. For time-lapse induction assays, induced cultures, prepared as described above, were immediately transferred after AHL inoculation to a CytoCapture imaging dish, incubated for 30 min at 37*°*C, and time-lapse imaged for 1 h.

### Competition assays with trans-cinnamaldehyde

Pure grade *trans*-cinnamaldehyde (CA, Sigma-Aldrich) was dissolved in absolute ethanol to a concentration of 5×10^−2^ M and further sub-diluted in water to concentrations ranging from 2×10^−2^ to 2×10^−3^ M. A flask with 10mL of LB, supplemented with 200 µg/mL ampicillin, was inoculated with a single colony from a freshly streaked plate of the *E. coli* biosensor strain. After incubation for 18 h at 37*°*C with vigorous shaking, the culture was diluted by a factor of 10^−3^ into 20 mL of supplemented M9 minimal medium and grown to an OD600 of 0.05 – 0.1 (∼4–5 h). 10µL of the CA solutions were mixed with 10µL of 1×10^−8^ M AHL _*fisch*_ and transferred to the wells of a flat-bottomed 96-well plate. 180-µL aliquots of the bacterial culture were added to each well, thus yielding an AHL _*fisch*_ final concentration of 5×10^−10^ M and CA concentrations ranging from 1×10^−4^ to 1×10^−3^ M. Preparation of blank and control wells, as well as incubation and assay conditions, have been described above. Growth rates were estimated from the slopes of linearly fitted plots of OD600 *vs.* time during the first 100 min of growth.

### Percolation function fitting

In its classical conception, PT describes the size of the cluster which is formed around the critical percolation threshold (Eq. 1). To test if the formation of the QS response in the biosensor could be explained by percolation kinetics, we fitted the experimental data in the region comprising the onset of the density-normalized response and before reaching the plateau to the percolation scaling law function:

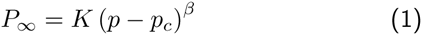

where the exponent *β* characterizes the abrupt rising of the order parameter in a phase transition system [60]. The value of the critical exponent solely depends on the dimensionality *n* of the lattice, thus making it universal. Initial attempts to mathematically reproduce the behavior of the biosensor, modeling the response as an infinite cluster of bacteria forming in a 3D lattice failed to yield the expected exponents for this dimensionality (not shown). By adding extra variables to account for the number of bacteria, *b*; the signal concentration, *s*; and time, *t*; we arrived at the following six-dimensional percolation equation:

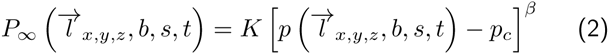

where 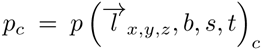 is the critical occupancy in the percolation lattice. Since we assumed that the AHL concentration remains constant during the experiment, Eq. 2 is simplified to a five-dimensional function of *P_∞_*:

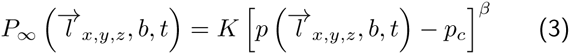

In simple percolation processes described in a five-dimensional lattice, *β* takes the universal value of 0.83 [32].

To test if the formation of the QS response in the biosensor could be explained by percolation kinetics, we fitted the experimental data in the region comprising the onset of the density-normalized response and before reaching the plateau to the percolation scaling law function (Eq. 1). This equation is equivalent to the three-parameter power Belehradek function (OriginLab, Northampton, MA), where *P_∞_* is the measured QS response (in our case violacein accumulation or FI/OD600), *K* is a proportionality constant, *t* is time, and *p_c_* becomes *t*_*c*_, *i.e.* is the critical time at which the response diverges and *β* is the critical percolation exponent [32]. The use of the Belehradek function to approximate percolation phenomena has been described [61, 62].

### Percolation analysis of published data

The plots of interest were captured as screenshots from the original papers (Table S1 summarizes the relevant information available in the original publications). The resulting images were uploaded to the free online application “web plot digitizer” (http://arohatgi.info/WebPlotDigitizer/app/). The axes were calibrated by selecting two known values for x and y, and indicating whether the axis was in log scale, when required. Data points on the original plot were selected on the image. The application returns the converted data as a text file. When the y-axis was specified as being in log scale, the application returned linearized data. The converted data was plotted and fitted to the percolation function as previously described. When possible, response data were normalized to cell density before fitting to the percolation function. In the case of Figure 4C, only A_480_ values higher than 0.4 in Figure 9 of Gallardo *et al.* [28] were considered for percolation analysis. These values were taken as the no-violacein baseline to discriminate violacein accumulation from absorbance due to cell growth of wt *C. violaceum*. For bioluminescence studies displaying the characteristic pit-like decrease of luciferase activity before the exponential rise (Figures 6B, C, and D) [36–38], the pit data points were excluded from the fitting, despite the fact that their presence has no effect on the percolation analysis (compare the two plots shown in Figure 6C, Table S1).

### Simulation of the AHL-based response of bacteria in a 3D-cubic lattice model

The AHL-based bacterial response was simulated in an artificial 3D-cubic lattice in which bacteria were assumed to live, move, and replicate in the presence of diffusing AHL and antagonist molecules, hereon named ligands (Figure S6A). The lattice was divided into *n*^3^ lattice wells, where *n* is the length of the lattice, and each of the wells have six neighboring positions connected by their edges. The lattice was programmed in networkx (1.8.1) [63]. Additional libraries were numpy (1.8.1) and scipy (0.13.3). The graphical user interface required guiqwt (2.3.2) and PyQt (4.9.6–1).

A complete pseudo-code representation of the simulation code is shown in Figure S6B. The complete open source code in Python, together with the stand-alone program, are available online (http://www.wwu.de/Biologie.IBBP/aggoycoolea/software/). The parameters driving bacterial growth, movement, and ligand diffusion, as well as the internal equilibrium kinetics, were calculated at discrete arbitrary time units for every bacterium and ligands present in the lattice. Bacteria are randomly seeded with defined starting densities, representing the fraction of spots that are occupied at the beginning of the simulation. Bacteria were allowed to move randomly along the edges as long as neighboring lattice cells were not occupied by other bacteria. The growth rate was defined as the probability of a given bacterium to duplicate itself in a unit of time whenever an adjacent lattice cell was available. Such growth rate, defining the fraction of bacteria that duplicated within the lattice ensured logistic growth, as described by Eq. 4, and resulted in growth curves that were in close agreement with experimental data (see Results section).

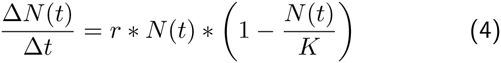

where *N* is the number of bacteria, *t* is the time, *r* is the growth rate, and *K* is the size of the 3D lattice (*n*^3^).

Ligands were also allowed to diffuse randomly across the lattice, being possible for more than one ligand to be present in the same lattice cell at a given time. When a bacterium and one or more ligand molecules coincide in the same lattice cell, the latter were assumed to enter the bacterium, where they could bind to free activator monomers under user-assignable probabilities. The probability of encounter between an activator molecule and its potential ligands (signal or antagonist) is determined as follows. For every lattice cell the algorithm creates a dynamic and random list of ligand species at every time point. The ligand species that becomes available for binding to the activator monomer is randomly chosen from this list. Thus, the more abundant a ligand species is the more likely that it will be chosen for binding. Whether binding between the activator and the ligand occurs depends on the probabilities associated to their corresponding *k* constants. Once a ligand molecule is chosen and bound to an activator, it is permanently removed from the list. Chosen, but unbound ligands are temporarily removed from the list until the next time point. The same sequence of events is performed for the next available activator until all activator molecules present in the lattice cell are allowed to bind. The list is dynamic and regenerates at every time step, according to ligand movement and availability in the lattice cells. The user can also assign probabilities to events such as activator synthesis and degradation, AHL release, dimerization, and dimer dissociation. A complete set of the parameters used for the different simulations is available in Table S3. For the sake of simplicity, all the events following dimerization were summarized in a single step. The number of monomers and dimers was randomly and asymmetrically distributed among the daughter cells [19, 64]. On the contrary, the QS status of a given cell was equally distributed among the two daughter cells upon cell division.

The concordance of our simulations with Hill behavior was analyzed as follows. After plotting the accumulation of density-normalized dimers *vs.* time at various signal densities, we calculated the rates of dimer accumulation by using the slopes of the linear region of the plots. Fitting to the Hill function was performed as described above.

Most simulations were run on personal computers and large simulations with a higher number of repetitions, and varying sets of parameters were carried out at the Morfeus GRID (Westfälische Wilhelms-Universität Münster, Germany) with Condor [65]. Additional details on the simulations are provided in Figure S6.

## Acknowledgments

This work was supported by the FP7 IIF Marie Curie project entitled BioNanoSmart_DDS (Contract No. 221111), and by funds for the Consolidation and structuration of competitive research units (Competitive Reference Groups) (REF. 2010/18), from the Spain Institute of Health “Carlos III” (Strategic Health Action, Project FIS PSI14/00059) and “Xunta de Galicia” (Project Competitive Reference Groups, 2014/043-FEDER). CVS was supported by a pre-doctoral fellowship of the Xunta de Galicia and by a FPU fellowship of the “Ministerio de Educación y Ciencia” of Spain, by a research fellowship of the DAAD (Germany), and a research fellowship of the Fundación Pedro Barrié de la Maza (Spain). We acknowledge support from The Danish Agency for Science, Technology and Innovation, Denmark (FENAMI project 10–093456); the research leading to these results has also received funding from the European Union’s Seventh Framework Programme for research, technological development and demonstration under grant agreement no. 613931. XQ was recipient of a fellowship from China Scholarship Council. We thank Christopher Anderson and Mariana Leguia for providing the plasmid pSB1A3-BBa_T9002, Carlos Bustamante for his support during the optimization of the *E. coli* fluorescent biosensor, and Ana Otero and Manuel Romero for providing the strain CV026 of *C. violaceum* and technical assistance with *C. violaceum* plate assays. We are also grateful to Antje von Schaewen for the generous access to the Safire Tecan-F129013 Microplate Reader and to Catalina Sueiro López for her help with the microscopy set up.

